# Structure and dynamics of differential ligand binding in the human ρ-type GABA_A_ receptor

**DOI:** 10.1101/2023.06.16.545288

**Authors:** John Cowgill, Chen Fan, Nandan Haloi, Victor Tobiasson, Yuxuan Zhuang, Rebecca J. Howard, Erik Lindahl

## Abstract

The neurotransmitter γ-aminobutyric acid (GABA) drives critical inhibitory processes in and beyond the nervous system, partly via ionotropic type-A receptors (GABA_A_Rs). Pharmacological properties of ρ-type GABA_A_Rs are particularly distinctive, yet the structural basis for their specialization remains unclear. Here we present cryo-EM structures of a lipid-embedded human ρ1 GABA_A_R, including a partial intracellular domain, under apo, inhibited, and desensitized conditions. An apparent resting state, determined first in the absence of modulators, was recapitulated with the specific inhibitor (1,2,5,6-tetrahydropyridin-4-yl)methylphosphinic acid and blocker picrotoxin, and provided a rationale for bicuculline insensitivity. Comparative structures, mutant recordings, and molecular simulations with and without GABA further explained the sensitized but slower activation of ρ1 relative to canonical subtypes. Combining GABA with picrotoxin also captured an apparent uncoupled intermediate state. This work reveals structural mechanisms of gating and modulation with applications to ρ-specific pharmaceutical design, and to our biophysical understanding of ligand-gated ion channels.

## INTRODUCTION

The neurotransmitter γ-aminobutyric acid (GABA) drives the majority of the inhibitory signaling in the mammalian nervous system via both ionotropic GABA_A_ and metabotropic GABA_B_ receptors. Differential pharmacology of GABA_A_ and GABA_B_ receptors has been critical in parsing their physiological functions using the specific inhibitors bicuculline (GABA_A_) or baclofen (GABA_B_) (Bormann, 1988; Hill and Bowery, 1981). An atypical class of GABA receptor was first identified from its insensitivity to both bicuculline and baclofen and was initially known as the GABA_C_ receptor (Drew et al., 1984). However, cloning and heterologous expression of these unusual receptors revealed that they are subtypes of the GABA_A_-receptor (GABA_A_R) family formed by ρ-type subunits (Cutting et al., 1991). Since their discovery, ρ-type GABA_A_Rs have been shown to play diverse roles in inhibitory signaling in neuronal and non-neuronal tissue. Their unique pharmacological profiles compared to canonical receptors have implications e.g. for differential bipolar cell signaling and visual transduction in the retina where it is highly expressed, (Dong et al., 1994; Matthews et al., 1994), neuroprotection (Yang et al., 2008, 2003), potential fear/anxiety responses in the lateral amygdala (Cunha et al., 2010), alcohol dependence (Blednov et al., 2014), sleep-wake cycles (Arnaud et al., 2001), and other physiological processes (Naffaa et al., 2017; Zhu et al., 2019).

Like canonical GABA_A_Rs, ρ-type GABA_A_Rs form pentameric ligand-gated ion channels (pLGICs) that pass chloride currents in response to binding GABA. They are composed of an N-terminal extracellular domain (ECD) followed in sequence by a transmembrane domain (TMD) of four transmembrane helices (M1–M4) per subunit. The neurotransmitter-binding site is formed at the subunit interface in the ECD, while the pore is lined by the second transmembrane helices (M2) of each subunit. Additionally, ρ-type GABA_A_Rs possess an intracellular domain between the M3 and M4 helices, also present though poorly conserved among other eukaryotic pLGICs.

The similarity of ρ-type receptors to canonical GABA_A_Rs extends little beyond homology and overall architecture. Whereas canonical GABA_A_Rs assemble as heteropentamers containing α, β and γ subunits (most commonly α1β2γ2), ρ-type GABA_A_Rs can form functional, GABA-activated homopentamers (Polenzani et al., 1991). These ρ-type GABA_A_Rs produce slowly activating and weakly desensitizing currents that are roughly 10 times more sensitive to GABA than the canonical receptor, and insensitive to classical modulators like barbiturates and benzodiazepines (Amin, 1999; Polenzani et al., 1991; Walters et al., 2000). The channel blocker picrotoxin (PTX) has been shown to be less effective at ρ1 versus other types of GABA_A_R, though the precise mechanism of open-versus resting-state block remains unclear for the GABA_A_R as a whole. The three ρ isoforms (ρ1-3) may also form heteropentameric assemblies with differing functional and pharmacological properties, though all known assemblies are inhibited by the specific ρ-type inhibitor (1,2,5,6-tetrahydropyridin-4-yl)methylphosphinic acid (TPMPA) (Murata et al., 1996).

Although structures of canonical GABA_A_R subtypes have been reported in recent years in apparent inhibited and desensitized states, the pharmacologically distinctive ρ-type receptor has been elusive. Currently available data also provide scant insight into GABA_A_R structure or dynamics in the absence of extracellular binding partners such as Fab fragments, or into the partially flexible intracellular domain, leaving open questions as to the structural conservation of gating and modulation mechanisms in this physiologically critical protein family. To address these gaps, here we report cryogenic electron microscopy (cryo-EM) structures and molecular dynamics simulations of a functionally validated construct of the human ρ1 GABA_A_R under apo, desensitized, inhibited, and blocked conditions in lipid nanodiscs.

## RESULTS

### Structure of human ρ1 GABA_A_Rs in resting, inhibited, blocked, and desensitized states

To gain insight into the structural basis for the unique pharmacological properties of ρ1 GABA_A_Rs, we solved cryo-EM structures in the absence of ligands and in the presence of TPMPA, PTX, and/or GABA to resolutions down to 2.2 Å (Figures 1A, S1 and S2, Table). Efforts to preserve the full-length sequence of human ρ1 GABA_A_R in HEK cells resulted in low expression and poor biochemical stability. Structural studies of other GABA_A_R subtypes often require removal of the intracellular domain in the M3–M4 loop to improve stability and expression (Miller and Aricescu, 2014; Zhu et al., 2018; Phulera et al., 2018; Kim et al., 2020). Using an AlphaFold2 model of the ρ1 GABA_A_R monomer to inform modifications, we removed regions predicted with low confidence and inserted a twin-Strep affinity-purification tag and thermostable GFP variant at the N-terminus and M3–M4 loop, respectively (Figure S2). This ρ1-EM GABA_A_R construct retained classical functional features of ρ wild-type (WT) GABA_A_Rs, including relatively slow gating kinetics and high affinity GABA activation with little desensitization (Figure S3A–B). These modifications substantially boosted expression and stability, resulting in a construct that could be stably purified and reconstituted into saposin nanodiscs for structural studies even in the absence of GABA (Figure S3C–D).

**Figure 1.**
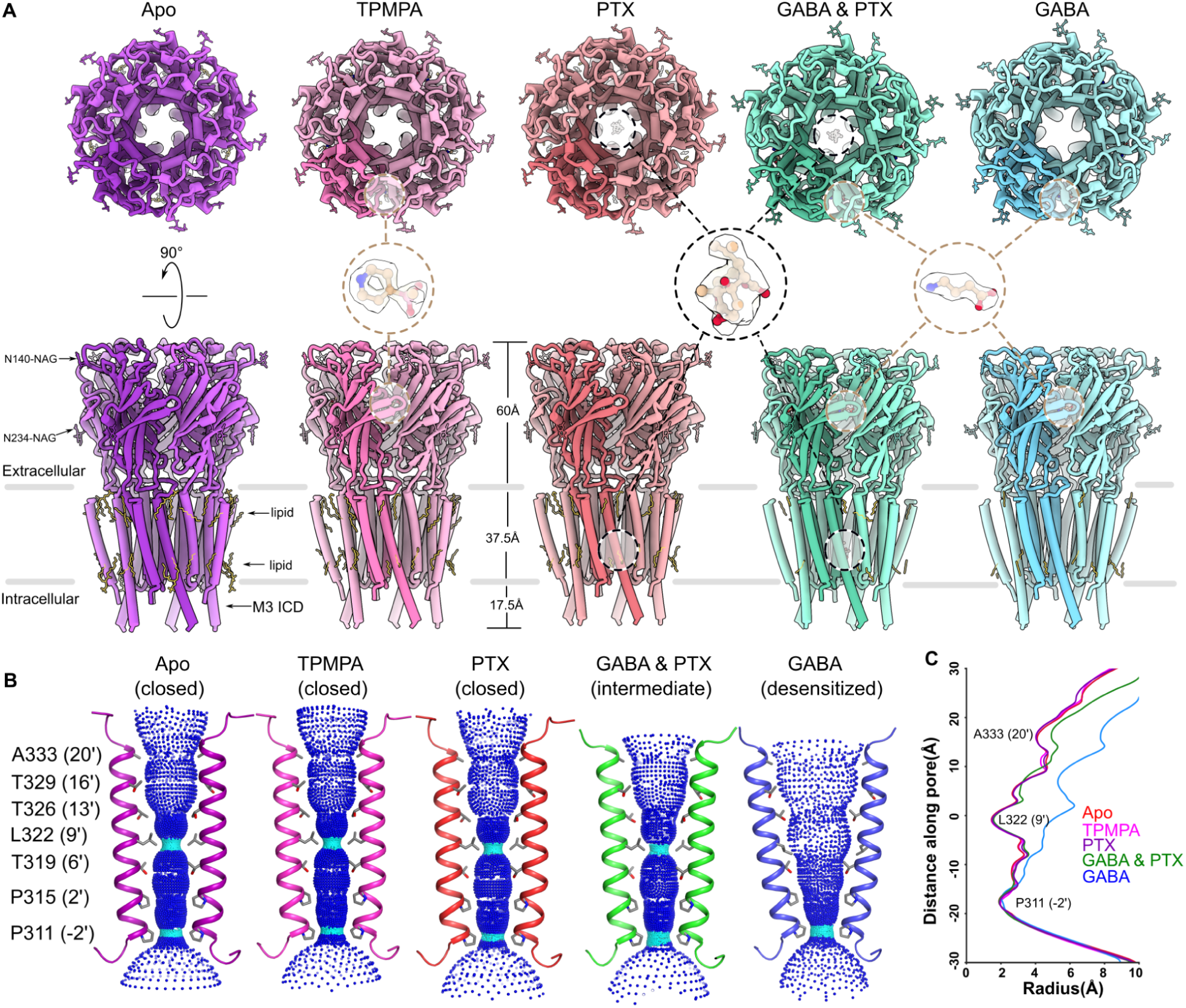
Structures of human ρ1 GABA_A_Rs in resting, inhibited, blocked, and desensitized states. (A) Cryo-EM structures of ρ1 GABA_A_Rs determined in the absence of ligand (apo, purple) or in the presence of TPMPA (pink), PTX (red), GABA plus PTX (green), or GABA (cyan), viewed from the extracellular side (top) or from the membrane plane (bottom). In each structure, a single subunit of the pentamer is colored darker for better definition. Ligands, glycans (NAG), and lipids are shown as sticks. Insets show TPMPA, PTX, and GABA with corresponding ligand densities in a representative orthosteric (brown dashed lines) or pore site (black dashed lines); oxygen, nitrogen and phosphorus atoms are colored red, blue and yellow respectively (by heteroatom). (B) Ion permeation pathways in ρ1 GABA_A_R cryo-EM structures, colored as in *A*. For clarity, only the M2 helices of two opposing subunits are shown as ribbons; amino acid residues facing the pore are shown as sticks, and labeled by residue number and prime notation at left. The pore radius of each structure was calculated using HOLE and displayed as dots. Cyan regions indicate constrictions to 1.15–2.3 Å (Smart et al., 1996). (C) Pore profiles of ρ1 GABA_A_R cryo-EM structures, colored as in *A*, with the Cα atom of L322 (9’) set to 0 along the y-axis.

Density maps enabled unambiguous assignment of most protein and ligand atoms, including features such as the cavity in the center of the tetrahydropyridyl ring of TPMPA and clear side-chain rotamers throughout the membrane-spanning regions (Figures 1A, S4A–B). Resolved protein regions also included a nearly three-turn extension of M3 in the intracellular domain beyond that of previous GABA_A_R structures (Figures S3E, S4B). This highly charged region of the protein (Figures S3F, S4C) showed only subtle differences across functional states, consistent with its proposed role interacting with scaffolding proteins rather than regulating channel gating (Jansen et al., 2008). In addition to protein, several additional densities in the transmembrane region were modeled as partial phospholipids (Figure S3G), including outer- and inner-leaflet sites previously identified in nicotinic acetylcholine receptors (Ananchenko et al., 2022). We also resolved two peripheral N-linked glycosylation sites (N140 and N234) per subunit (Figure S4D), with no sign of the vestibular glycosylation observed in α subunits (Kim and Hibbs, 2021; Phulera et al., 2018; Zhu et al., 2018). The N140 glycosylation site was located at the upper ECD β2–β3 loop, similar to one of the sites in β subunits (N80 in β3), and possibly conserved in ρ2 based on sequence alignment (Figure S2). The N234 glycosylation site was located in helix 5 N-terminal to loop F, and constituted a unique feature of ρ1.

Based on the radius of the P311 (-2’) and L322 (9’) gates (where primes denote positions on M2 relative to a conserved basic residue at the cytosolic end), the apo, TPMPA, and PTX-bound structures appear to represent resting states of the pore. The GABA-bound structure is likely to be desensitized; the pore radius at -2’ (2.0 Å) is slightlylarger than that observed for canonical receptors (≤1.8 Å) (Kim et al., 2020; Laverty et al., 2019; Masiulis et al., 2019), but still not expanded enough to conduct at least fully hydrated chloride ions (3.2Å). (Figure 1B–C). The structure with GABA as well as PTX (GABA & PTX) adopted an intermediate state, with the pore similar to the resting state, but the ECD tilted and rotated in a similar way as in the desensitized state as described in detail below (Video S1).

### TPMPA inhibits channel gating via conformational selection of a distinctive resting state

Existing structures thought to represent resting states of GABA_A_Rs have included competitive inhibitors, allosteric inhibitors, or serendipitous density in the orthosteric ligand-binding site (Kim et al., 2020; Laverty et al., 2019; Masiulis et al., 2019). Studies of heteromeric receptors have also relied on fiduciary nanobodies or megabodies, which aid subunit classification but are known to modulate channel gating (Laverty et al., 2019; Masiulis et al., 2019; Phulera et al., 2018; Zhu et al., 2018) (Figure S3H–I). The stability and symmetry of the ρ1-EM GABA_A_R construct (described hereafter for simplicity as ρ1) enabled us to determine the structure in the absence of any additives, making this the only unliganded GABA_A_R structure yet reported, to the best of our knowledge. Compared to apo conditions, both the global structure of the receptor and local geometry at the neurotransmitter-binding site showed striking similarity in the presence of the ρ-type GABA_A_R-specific inhibitor TPMPA (Figure 2A–B). Even residues directly involved in coordination (e.g. F159 on loop A; E217 and Y219 on loop B; Y262, S264, Y268 on loop C; and R125 on loop D of the complementary subunit, denoted here (-)) underwent minimal rearrangements upon inhibitor binding (Figure 2B–C). Thus, we posit that TPMPA acts by conformational selection of the resting state, rather than an induced-fit mechanism. Pore profiles of ρ1 in both resting-state structures were largely comparable to canonical GABA_A_Rs in the presence of bicuculline and to apo glycine receptors (GlyRs), with a near identical constriction at the 9’ gate, and a slightly expanded outer vestibule (Kim et al., 2020; Kumar et al., 2020; Masiulis et al., 2019) (Figure S5A).

**Figure 2.**
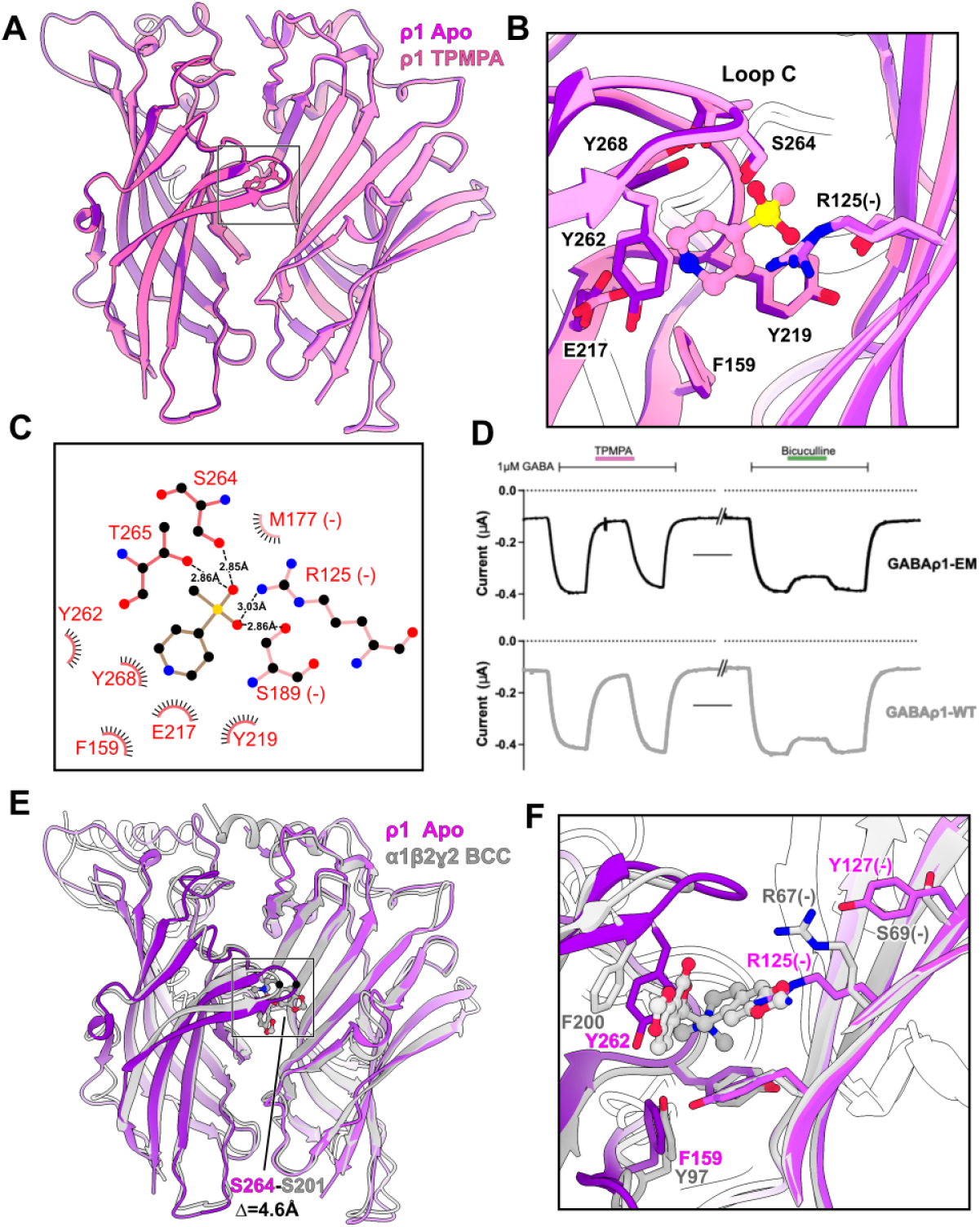
TPMPA inhibits via ρ1-specific selection of the resting state. (A) Superimposed ECDs of apo (purple) and TPMPA-bound (pink) ρ1 structures. For clarity, only two subunits are shown. Box indicates zoom region in *B–C*. (B) Zoom view of ligand-binding sites in apo and TPMPA structures, colored as in *A*, showing several residues interacting directly with TPMPA as sticks. The TPMPA molecule is shown as ball-and-stick. Side chains and TPMPA are colored by heteroatom. (C) Schematic of ρ1 interactions with TPMPA, created in LIGPLOT (Wallace et al., 1995). Electrostatic interactions are indicated as dashed lines to fully modeled side chains, and labeled by inter atomic distance. Hashes indicate hydrophobic interactions. (D) Representative current traces upon application of 2 μM TPMPA or 100 μM bicuculline during pulses of 1 μMGABA to oocytes expressing ρ1-EM (top, black) or -WT constructs (bottom, gray). Scale bars represent 50 seconds. (E) ECDs of apo ρ1 (purple) and the bicuculline (BCC)-bound α1β2γ2 GABA_A_R (gray, PDB ID 6X3S), superimposed on the complementary of two depicted subunits. Label indicates relative displacement of the outermost loop-C Cα atom in ρ1 (S264) versus that in β2 (S201). Box indicates zoom region in *F*. (F) Zoom view of ligand-binding sites in apo and BCC-boundα1β2γ2 GABA_A_Rs, colored as in *E*. Several key residues are shown as sticks; the BCC molecule is shown as ball-and-stick.

The bicuculline resistance that led to the discovery of ρ-type GABA_A_Rs was retained in our ρ1-EM GABA_A_R construct (Hill and Bowery, 1981) (Figures 2D, S5B), and could be rationalized by comparing our apo structure with bicuculline-bound structures of canonical neuronal GABA_A_Rs (Kim et al., 2020; Masiulis et al., 2019) (Figure 2E–F). Compared to these previous cases, Loop C in the ρ1 resting state adopted a more closed conformation over the orthosteric ligand site, resulting in a pocket too small to accommodate bicuculline. If superimposed into the resting ρ1 pocket, bicuculline would clash with residues Y262 from loop C and R125(-) from loop D. Although these residues are conserved at orthosteric β/α interfaces in canonical GABA_A_Rs, nearby variations appeared to explain their structural differences. Loop C was extended by one residue in ρ1 relative to other GABA_A_R subunits, such that Y262 displaced deeper into the ligand site (Figures S2, S3E, 2E–F). On loop D, the position of R125(-) appeared to be restricted to an orientation towards the ligand site by the bulky side chain of Y127(-), which in α1 subunits is replaced by a more accommodating serine. Thus an elongated loop C and restricted R125(-) snugly coordinated TPMPA while limiting accessibility to bicuculline. This result offers a structural rationale for previous chimera studies, in which constructs including substitutions in loop C and Y127 were shown to swap bicuculline sensitivity between ρ1 and α1β2γ2 GABA_A_Rs (Jianliang Zhang et al., 2008).

### Structural determinants of ρ-type high-sensitivity GABA activation

The structure of GABA-bound ρ1 offered a structural rationale for activation, including the higher sensitivity and slower kinetics of ρ-type versus canonical GABA_A_Rs (Enz and Cutting, 1998). Our desensitized structure shows GABA tightly coordinated at the intersubunit interface, with the amine group of the agonist in contact with an aromatic cage and conserved glutamate (E217) on the principal subunit, while the carboxylate could form a salt bridge with R125 on the complementary subunit (Figure 3A–B). The coordination of GABA in ρ1 resembles that of the canonical α1β2γ2 GABA_A_R, aside from the elongated ρ1 loop C (Figure 3C–D). Overlaying the neurotransmitter site in the apo and GABA-bound structures reveals surprisingly subtle local rearrangements upon GABA binding, with loop C closing over the site by around 2 Å (Figure 3E–F).

**Figure 3.**
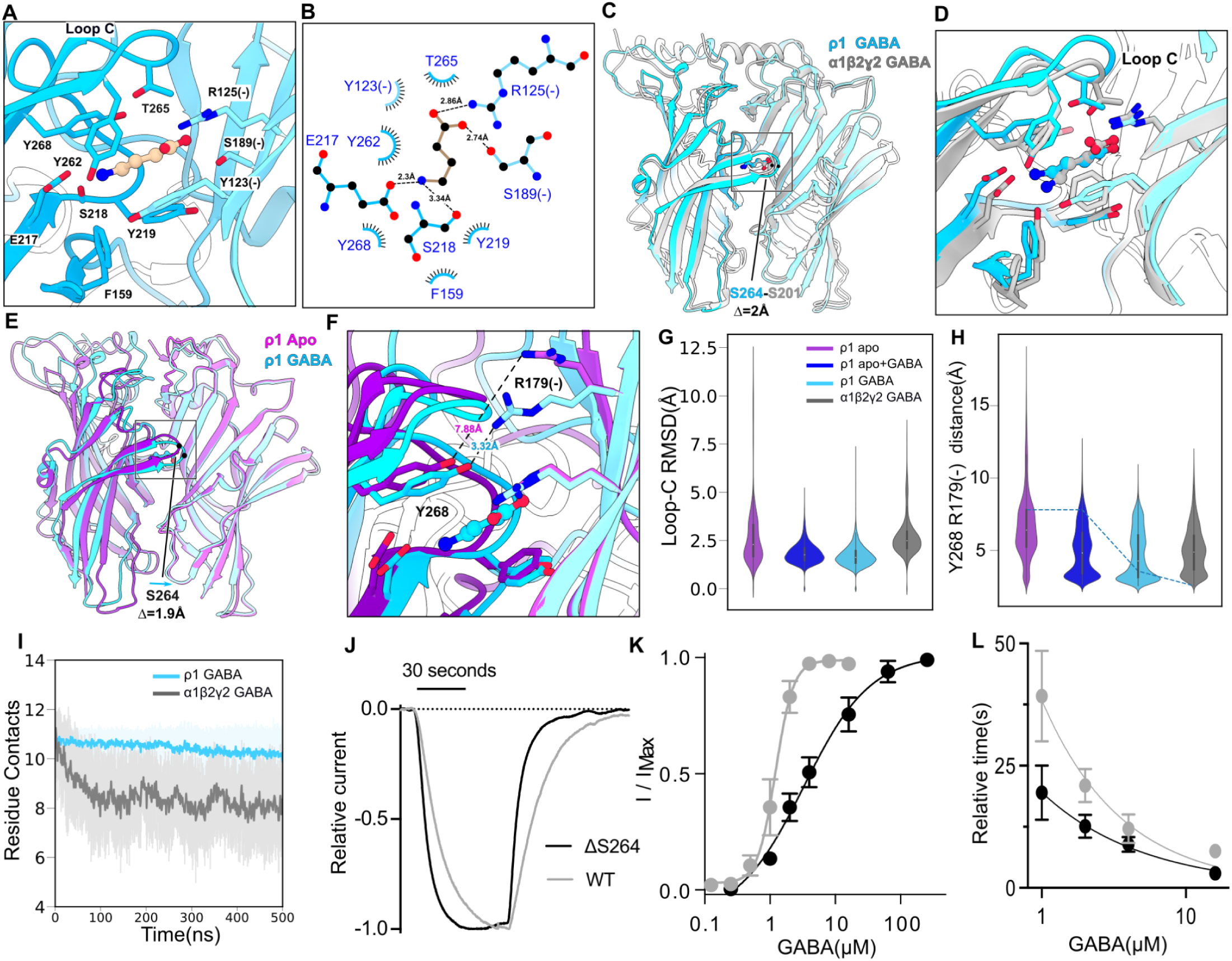
Structural determinants and binding-site closure associated with GABA sensitivity. (A) Zoom view of the neurotransmitter-binding site in GABA-bound ρ1. GABA (wheat) and its interacting side chains are colored by heteroatom and shown in ball-and-stick and stick representations, respectively. (B) LIGPLOT schematic of TPMPA interactions. Electrostatic interactions are indicated as dashed lines to fully modeled side chains, and labeled by interatomic distance. Hashes indicate hydrophobic and cation-π interactions. (C) ECDs of GABA-bound ρ1 (cyan) and α1β2γ2 GABA_A_Rs (gray, PDB ID 6X3Z), superimposed on the complementary of two depicted subunits. Label indicates relative displacement of the outermost loop-C Cα atom in ρ1 (S264) versus that in β2 (S201). Box indicates zoom region in *D*. (D) Zoom view of neurotransmitter-binding sites in GABA-bound ρ1 and α1β2γ2 GABA_A_Rs, colored as in *C*. (E) ECDs of apo (blue) and GABA-bound (purple) ρ1 structures, superimposed on the complementary of two depicted subunits. Label indicates relative displacement of the outermost loop-C Cα atom (S264). Box indicates zoom region in *F*. (F) Zoom view of neurotransmitter-binding sites in apo and GABA-bound ρ1, colored as in *E*. The nearest distance between heavy atoms of Y268 and R179(-) in each structure is labeled. (G) Violin plots of Cα RMSD for residues in loop C, averaged across all simulation replicates and subunits, calculated by aligning the ECD of each subunit independently to the starting structure. (H) Violin plots of average distance between the hydroxyl oxygen of Y268 (Y205 in β2) and amine nitrogen of R179(-) (R120 in α1), averaged across simulation replicates and relevant subunits. Initial distances for each system are depicted in dotted lines. (I) Number of amino acid residues within 3.5 Å of GABA during 500-ns molecular dynamics simulations, averaged over all simulation replicates and relevant subunits. (J) Sample traces showing activation of WT (gray) and ΔS264 (black) ρ1 constructs by 2 μM GABA, normalized to the steady-state current, showing accelerated activation and deactivation in the deletion mutant. (K) Concentration-response curves based on peak GABA activation of WT (gray) and ΔS264 (black) ρ1, showing decreased apparent affinity in the deletion mutant. Circles represent mean currents ± standard error, normalized to the maximal activation recorded for each construct (I/I_max_), *n* = 4 (WT) or 5 (ΔS264); solid lines indicate nonlinear regression fits. (L) Rise time (in seconds) of activation for WT (gray) and ΔS264 (black) ρ1 by various concentrations of GABA, showing relative acceleration in the deletion mutant. Circles represent mean 10-90% rise time ± standard error, *n* = 4; solid lines indicate exponential decay fits.

To elucidate the differential dynamic as well as structural effects of GABA binding, we carried out all-atom molecular simulations of the apo and desensitized ρ1 structures, as well as the GABA-bound α1β2γ2 GABA_A_R (Figure 3G–I). Despite comparable coordination in static structures, GABA remained more tightly bound to ρ1 versus α1β2γ2 subtypes, as indicated by reduced root-mean-square deviations (RMSD) and more consistent direct amino-acid contacts (Figures S5C–D, 3I). In particular, GABA made relatively transient contacts in α1β2γ2 versus ρ1 with aromatic-box residues F159 and Y262, as well as loop-C residue T265, which has been shown to be critical for GABA binding (Naffaa et al., 2016). Consistent with the static structures, loop C remained more tightly closed and sampled a narrower distribution in the desensitized ρ1 system than in the resting ρ1 or desensitized α1β2γ2 GABA_A_R systems (Figure 3G).

To capture initial transitions induced by GABA binding, we also simulated the apo ρ1 structure with GABA inserted into the binding site. GABA remained bound at all five sites of the homopentamer across 3 μs total simulation time, with comparable stability to the experimental structure of GABA-bound ρ1, and lesser deviations than in the GABA-bound α1β2γ2 subtype (Figures S5C). Moreover, adding GABA to the apo structure stabilized a tightened configuration of loop C, similar to the experimental GABA-bound ρ1 (Figure 3G). This configuration enables a hydrogen bond between loop-C residue Y268 and loop-F residue R179(-), evident in GABA-bound but not apo systems (Figure 3F,H). Interestingly, mutations of R179 in ρ1 GABA_A_Rs have been shown to eliminate GABA activation without disrupting trafficking to the plasma membrane (Harrison and Lummis, 2006). Thus, formation of this hydrogen bond is likely an early event in the activation pathway coupling GABA binding to pore opening.

The structures and simulations above indicate that a distinctive configuration of the orthosteric binding site, including an extended loop C, primes and stabilizes ρ1 for activation by GABA. Furthermore, this primed apo conformation could partially limit GABA access to the binding site, slowing activation. Indeed, radioligand binding assays have suggested that GABA only has limited access to its binding site in ρ1 (Chang and Weiss, 1999). To directly assess the role of the extended loop C in ρ1 gating, we recorded GABA activation upon deleting S264, the unaligned residue located at the tip of loop C (Figure S2). This mutation accelerated activation relative to the WT ρ1 receptor, while reducing the apparent GABA affinity (Figure 3J–L). This functional result further substantiates an important role for the elongated loop C of ρ1 both in enhancing GABA binding and slowing GABA entry into the binding site.

### Global rearrangements upon GABA binding

To understand the gating mechanism of ρ1, we compared its structure and dynamics under apo and GABA-bound conditions. Alignment of these structures on the TMD reveals that, in contrast to the modest local rearrangements detailed above, GABA binding induces more dramatic global conformational changes (Figure 4). In the GABA-bound versus apo structures, the ECD β-sandwich of the principal subunit has tilted towards that of the complementary subunit, translocating the outermost ECD 9.3 Å and the innermost tip 2.9 Å; loop C in particular is displaced 7.2 Å (distances between Cɑ atoms of G93, S203, or S264 residues in neighboring subunits, Figure 4B–C). Bridged by GABA interactions, the ECD interface between adjacent subunits is also more compact, with a 110-Å^2^ (10%) increase in the total buried surface area between two subunit ECDs. In MD simulations of the same structures, GABA binding was associated with general contraction of the outermost ECD (Figure 4H), and also of individual β-sandwiches facing the domain interface (Figure 4I). GABA binding also induces a relative rotation of the ECD 10° counterclockwise when viewed from the extracellular space, a twist which is also preserved in our MD simulations (Figure 4J). Adding GABA to the apo structure was not sufficient to induce domain rotation nor contraction of the outer or inner ECD within simulation timescales (500 ns; Figure 4H–J), indicating these transitions proceed later in the gating cycle than the loop-C effects described above.

**Figure 4.**
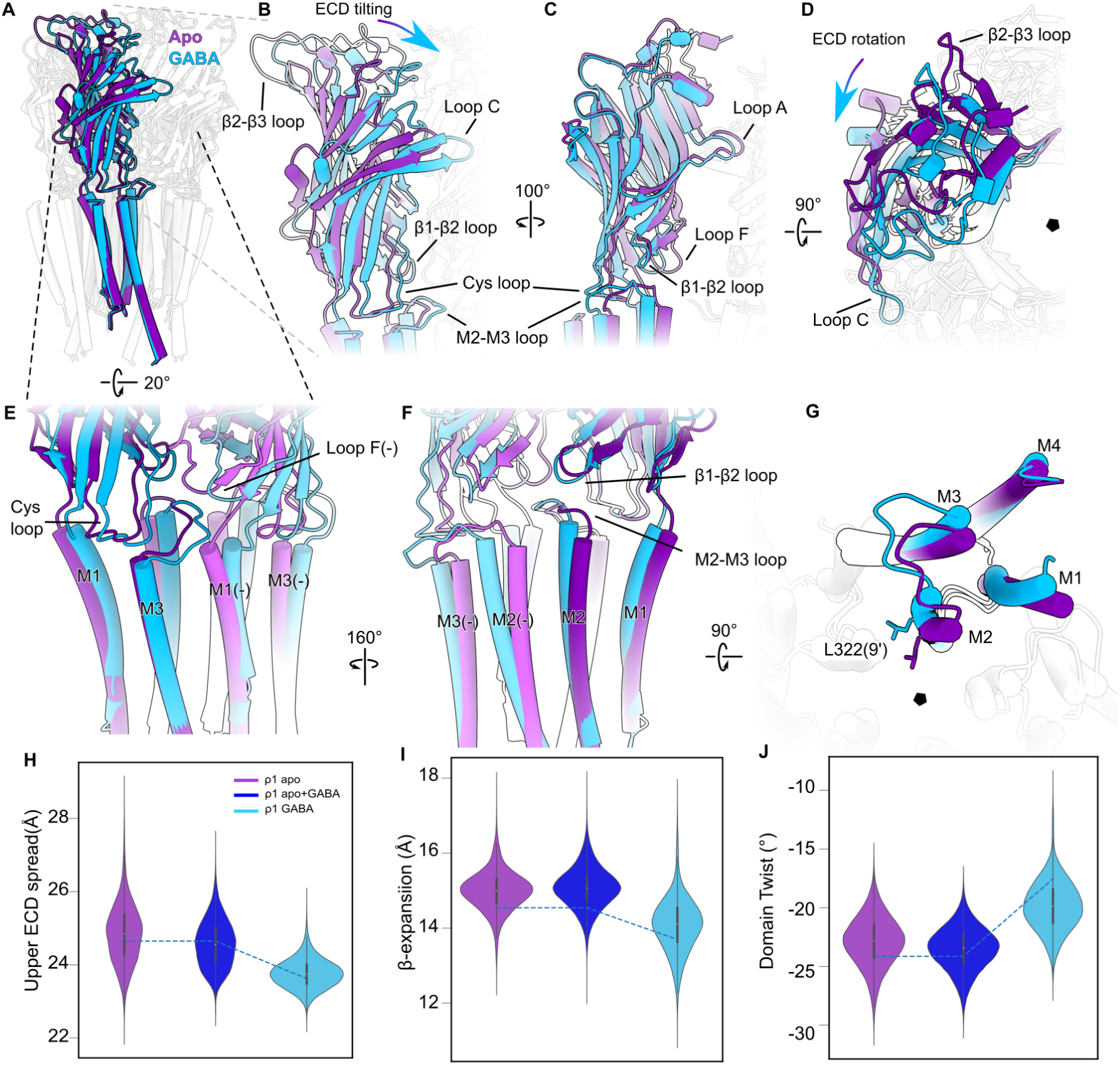
Global rearrangements upon GABA binding. (A) Superimposed apo (purple) and GABA-bound (cyan) ρ1 structures, viewed from the membrane plane and aligned on the TMD. For clarity, all but one subunit are mostly transparent. Gray and black dashed lines indicate zoom regions in *B–D* and *E–G*, respectively. (B) Zoom view of the ECD, represented as in *A*, showing rigid-body tilting upon GABA binding. (C) Zoom view of the ECD as in *B*, rotated by 100° for a view from the outer vestibule. (D) Zoom view of the ECD as in *B–C*, rotated by 90° for a view from the extracellular side, showing domain rotation upon GABA binding. (E) Zoom view of the ECD-TMD interface, represented as in *A*, showing loops translation upon GABA binding. (F) Zoom view of the ECD-TMD interface as in *E*, rotated by 160° for a view from the channel pore. (G) Zoom view of a single subunit TMD as in *E–F*, rotated by 90° for a view from the extracellular side, showing helix and loop movements upon GABA binding. The hydrophobic gate at L322 (9’) is shown as sticks; black pentagon indicates the pore axis. (H) Violin plots of global spread in the outward-facing ECD (upper ECD spread, Å) during simulations of the apo ρ1 structure in the absence (purple) and presence (blue) of GABA, and of the GABA-bound ρ1 structure (cyan). Each plot represents four replicate 500-ns MD simulations, showing relative contraction in the GABA-bound structure. Dotted lines indicate initial values calculated from the experimental structure. (I) Violin plots for β-sandwich spread at the domain interface (β-expansion, Å) during simulations, represented as in *H*, showing relative contraction in the GABA-bound structure. (J) Violin plots for relative rotation of the ECD versus TMD (domain twist, °) during simulations, represented as in *H–I*, showing twisting towards less negative values in the GABA-bound structure.

These tilting, contracting, and rotating motions of the ECD are transmitted to the TMD via the domain interface, mediated both by the covalent linkage in pre-M1 and noncovalently through interactions between the β1–β2 loop, Cys loop, loop F(-), and the M2–M3 loop. Specifically, the Cys loop and loop F move counter-clockwise toward the complementary subunit core, while the β1–β2 loop displaces down toward the TMD (Figure 4E–F). The M2–M3 loop is sandwiched between these ECD motifs, such that it also translocates toward the complementary subunit and away from the pore axis (Figure 4E–G). These movements are mediated in part through nonpolar interdomain interactions: upon GABA binding, V114 in β1–β2 and F205 in the Cys loop form hydrophobic contacts with M2–M3 residues P336 and M335, respectively (Figure S6). An electrostatic triad is maintained in resting and desensitized states between R279 in pre-M1, E113 in β1–β2, and D208 in the Cys loop, all largely conserved residues that have been implicated in stabilizing the domain interface (Kash et al., 2003; Price et al., 2007; Xiu et al., 2005). In the desensitized state, E113 can also form an intersubunit salt bridge with M2–M3 residue R337 (Figure S6C), at which mutations have been shown to affect channel activity (Xiu et al., 2005).

Structural rearrangements at the domain interface further propagate to the TMD, tilting the outward-facing end of each transmembrane subunit away from the pore axis, including a 4.3-Å expansion of the cleft between neighboring M2 helices (distance between N332 residues, 19’) (Figure 4E–G). These shifts propagate to the hydrophobic gate formed by the L322 (9’) residues, which shift and rotate away from the channel pore towards the subunit interfaces, expanding the presumed activation gate. In place of the 9’ constriction, the narrowest point in the pore of the GABA-bound structure is at the -2’ gate, which at 2.0 Å is still too narrow to pass hydrated chloride ions. The overall profile of this apparent desensitized state resembles that of homomeric β3 GABA_A_Rs and α1 glycine receptors; interestingly, the 9’ activation gate is 4 Å wider than in GABA-bound structures of α1β2γ2 and α1β3γ2 GABA_A_Rs, although this profile varied more extensively during MD simulations of the desensitized versus resting states of ρ1 (Figure S5E–F).

Tilting of the transmembrane helices upon GABA binding weakens interactions across the outward-facing intersubunit interface, expanding the cleft between the M3 and M1(-) helices on neighboring subunits. A similar expansion of the intersubunit TMD interface was previously observed in canonical GABA_A_Rs, facilitating binding of positive allosteric modulators such as barbiturates and benzodiazepines (Kim et al., 2020). However, ρ-type GABA_A_Rs are notably insensitive to pharmaceuticals that act at this allosteric site. In all of our ρ1 structures, the outward intersubunit cleft is occluded by residues including I328 (15’) and W349 on M2 and M3 respectively (Figure S6D). Mutations at these residues have been shown to sensitize ρ1 to both pentobarbital and diazepam (Amin, 1999; Belelli et al., 1999; Walters et al., 2000), supporting a critical role for steric accessibility at this site in subtype-specific drug effects.

### PTX uncouples ECD movements from channel gating

Initial studies on native ρ-type GABA_A_Rs from retina indicated low sensitivity to the plant alkaloid PTX (Feigenspan et al., 1993) relative to canonical subtypes. However, subsequent studies demonstrated PTX blocks ρ1 currents at ∼10 times higher doses, and with stronger dependency on GABA concentration (Qian et al., 2005, 1998; Wang et al., 1995). Our structures show that PTX binds in both the presence and absence of GABA within the channel pore, situated between P319 (2’) and the 9’ activation gate (Figure 5A–C). This site and pose are similar to those reported for the α1β3γ2 GABA_A_R, though resolution was not sufficient to exclude at least partial occupancy of an inverted pose observed in the α1β2γ2 GABA_A_R (Kim et al., 2020; Masiulis et al., 2019). Interestingly, the proline at 2’ is specific to ρ1 GABA_A_Rs (Figure S2), and mutation of this position to amino acids found at the equivalent position in other GABA_A_Rs (including ρ2) increase apparent PTX affinity 10-fold while reducing the competitive component of PTX action on GABA binding (Wang et al., 1995).

**Figure 5.**
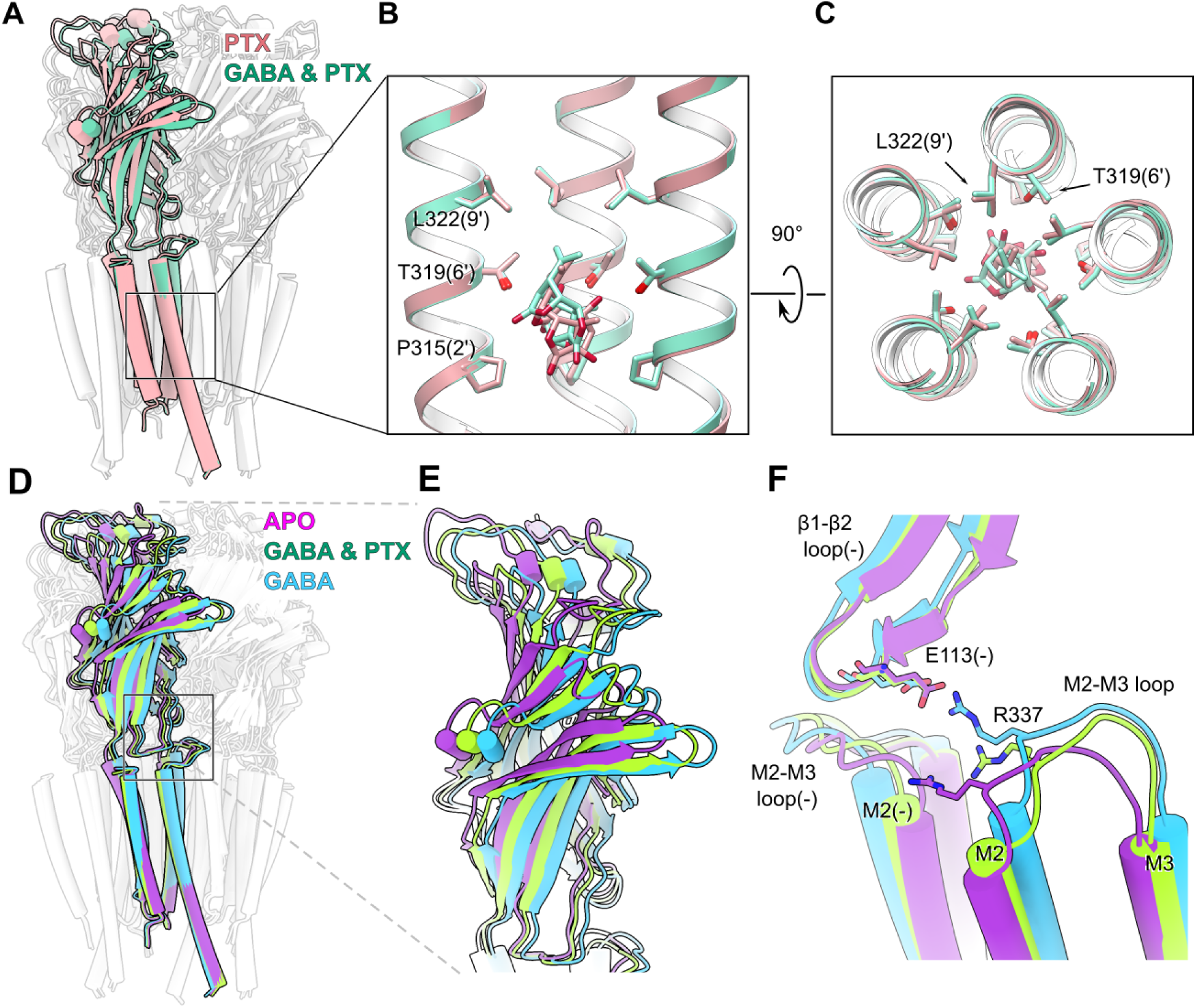
PTX uncouples ECD movements from channel gating. (A) Superimposed ρ1 structures with PTX, in the absence (red) or presence (green) of GABA, aligned on the TMD and viewed from the membrane plane. For clarity, all but one subunit is mostly transparent. Box indicates zoom region in *B–C*. (B) Zoom view of the inner M2 helices surrounding the PTX site, represented as in *A*. For clarity, only the three most distal subunits are shown, providing a view from the pore interior. PTX and the pore-lining residues it contacts (9’, 6’, and 2’) are shown as sticks and colored by heteroatom. (C) Zoom view of all five M2 helices, represented as in *B*, rotated by 90° for a view from the extracellular side. (D) Superimposed apo (purple), GABA & PTX (green) and GABA-bound (cyan) ρ1 structures, represented as in *A*. Dashed lines and solid box indicate zoom regions in *E* and *F*, respectively. (E) Zoom view of a single subunit of the ECD, represented as in *D*. (F) Zoom view of the intersubunit R337-E113(-) interaction between the M2–M3 and β1–β2 loops, represented as in *E*.

Binding of PTX alone was associated with little change in the overall structure relative to the apo condition (Figure S7A). Conversely, binding of PTX in the presence of GABA resulted in a structure with mixed characteristics (Figures 5D–E, S7B). The ECD of the GABA & PTX structure largely resembles the GABA-bound desensitized structure, while the TMD more closely resembles resting-state structures. Indeed, superimposing PTX into the desensitized state of ρ1 showed a decrease in contacts with pore-lining residues, indicating the toxin binding is more favorable in a resting-like pore, which could thus be stabilized (Figure S7C). This profile is generally consistent with previous structures of canonical GABA_A_Rs, and distinct from the apparent open state of PTX-bound GlyR (Kim et al., 2020; Kumar et al., 2020; Masiulis et al., 2019) (Figure S5G). The domain interface of the GABA & PTX structure is distinct from both resting and desensitized states, characterized by an intermediate configuration of the outer M2–M3 helices, especially notable in the intersubunit R337-E113(-) interaction between the M2–M3 and β1–β2 loops (Figures 5F, S6B). Additionally, in contrast to the desensitized GABA-bound structure, the ECD shows a lesser degree of rotation relative to the TMD for the GABA+PTX system (Figure 5D–E). Thus, we posit this structure describes an intermediate state, trapped between resting and desensitized states (Video 1).

Previous structures of α1β2γ2 and α1β3γ2 GABA_A_Rs with PTX disagree regarding the mechanism of PTX block (Kim et al., 2020; Masiulis et al., 2019). Structures of α1β3γ2 suggest a blocking mechanism specific to the resting-like pore, while the α1β2γ2 structure, as well as functional experiments, suggest that PTX binds to a partially open or desensitized pore. In light of this discrepancy, we built an allosteric model of PTX action that can account for both the structural and functional effects of PTX we observe on ρ1 GABA_A_Rs (Goldschen-Ohm et al., 2014) (Figure S7D). According to this model, PTX can bind to both the closed and open pore in the presence as well as absence of GABA. State dependence of PTX binding arises from the coupling energy to the activation gate, resulting in 3.2 kcal/mol more favorable binding to the resting rather than open gates. At saturating concentrations of GABA and PTX, the PTX binding would stabilize the resting pore, which then exceeds the coupling energy supplied by GABA binding, and prevents opening of the activation gate. Consistent with this model, in electrophysiological recordings of WT ρ1 we observed a rightward shift in the apparent affinity and a reduction in the efficacy of GABA at increasing concentrations of PTX (Figure S7E). This effect is also consistent with the uncoupling of ECD and TMD movements observed in the GABA & PTX structure. Mutations of the unique 2’ proline in ρ1 would be expected to alter state-dependent stabilization of PTX binding and thus the coupling energy between PTX binding and activation gating, helping to reconcile how local changes in sequence at this position can eliminate the competitive component of picrotoxin block in ρ1 GABA_A_Rs (Wang et al., 1995).

## DISCUSSION

The structures, simulations, and functional experiments above substantiate a mechanistic model for gating in ρ1 GABA_A_Rs. According to this scheme, the resting state—captured first as a genuine apo structure, without modulators or binding partners—can be selectively stabilized by a competitive antagonist (TPMPA) or the blocker PTX (Figure 6A, left). As demonstrated both in static structures and MD simulations, agonist binding initiates rapid closure of loop C over the orthosteric site, followed by longer-range contraction and rotation of the ECD. In the absence of blocker, the resulting conformational wave leads to expansion of the outer transmembrane pore to enable chloride conduction, followed by contraction of the inner gate to produce a desensitized state (Figure 6A, right). In the presence of PTX, GABA-induced conformational changes in the ECD are reduced, and moreover uncoupled from those in the TMD, capturing a presumed intermediate (Figure 6A, center). This mechanism is broadly similar to those proposed for other pLGIC, helping to assuage concerns that fiduciaries known to modulate gating, or inhibitors used to trap resting-state structures, may substantially alter the conformational landscape (Video S2, Kim et al., 2020; Kumar et al., 2020; Masiulis et al., 2019; Noviello et al., 2021; Yu et al., 2021).

**Figure 6.**
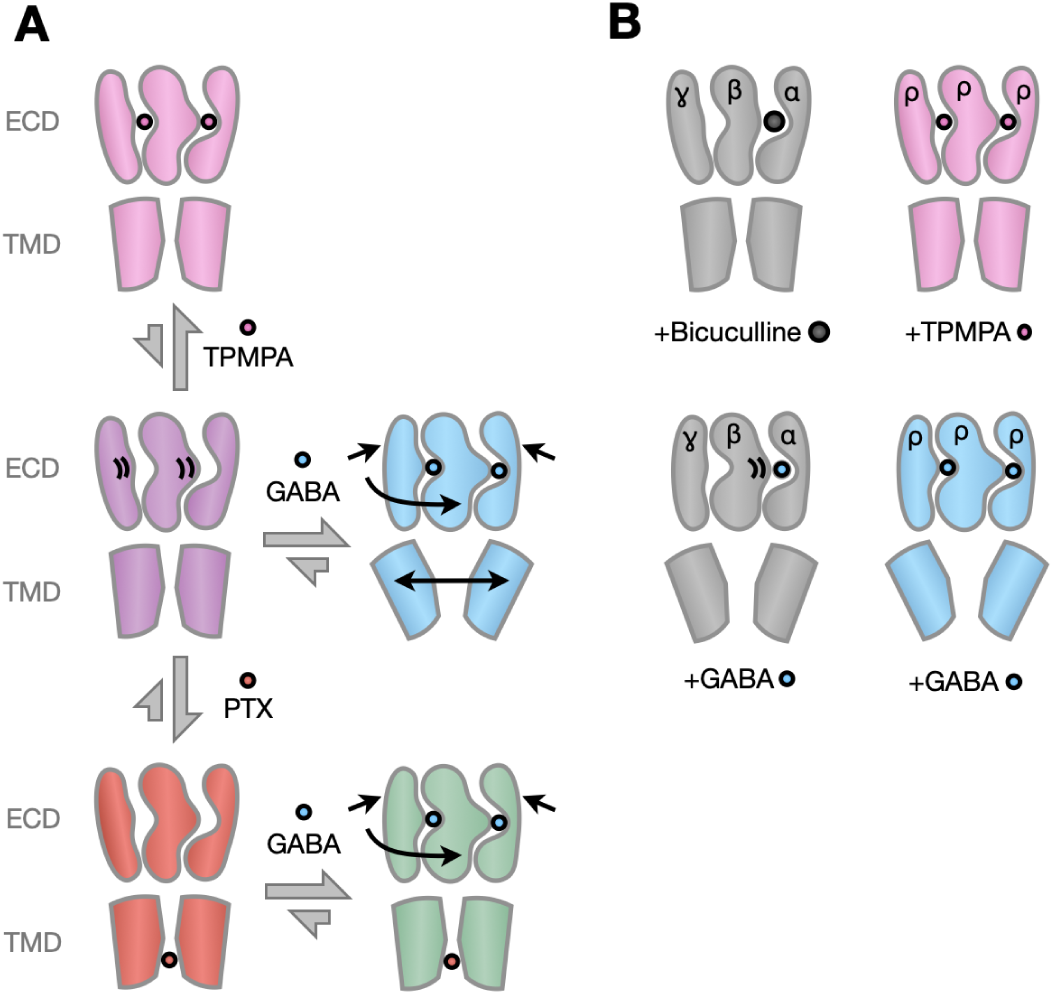
Proposed mechanism and differential features gating in ρ GABA_A_Rs. (A) In the proposed ρ-GABA_A_R conformational cycle, the resting state (purple) can be preferentially stabilized by binding TPMPA in the ECD (pink) or PTX in the pore (red). Binding GABA initiates closure of loop C over the orthosteric binding site, along with longer-range contraction and rotation of the ECD; in the absence of blocker, the resulting conformational wave leads to expansion of the outer transmembrane pore to produce an open or desensitized state (cyan). In the presence of PTX, GABA-induced conformational changes in the ECD are uncoupled from those in the TMD, capturing a presumed intermediate (green). Gray arrows indicate conformational equilibria; black arrows indicate structural changes in the labeled model relative to its left hand precursor. For clarity, each model shows three (of five) neighboring subunits in the ECD, and two opposing subunits in the TMD. (B) Cartoons as in *A*, showing that extension of loop C in ρ versus αβɣ GABA_A_Rs prefers binding of TPMPA over the larger inhibitor bicuculline (top), and facilitates stable binding of GABA (bottom). Colored models correspond to ρ1 structures reported in this work; gray models were previously reported for canonical neuronal subtypes, as described in the text.

The only well resolved class in our GABA dataset contained a constricted inner gate, smaller than a fully hydrated chloride ion, indicating a desensitized state (Figure 6A, right). This result was initially surprising, as fast desensitization is limited in ρ1, including our ρ1-EM construct (Figure S3A). However, the maximal open probability of ρ-type GABA_A_Rs in saturating GABA is unclear, due in part to their low unitary conductance (Chang and Weiss, 1999; Wotring et al., 1999). Estimates based on GABA binding and ionic currents in single oocytes indicate that only 10% of available receptors may be conductive at a given time (Chang and Weiss, 1999; Colquhoun, 1999). Thus, the fraction of open receptors may be insufficient to identify during 3D classification. Still, we cannot rule out the possibility that the choice of lipids, nanodisc scaffold, or other experimental conditions influenced the conformational equilibrium of ρ1, as was recently observed for the prokaryotic channel ELIC (Petroff et al., 2022; Vikram Dalal et al., 2022). On the other hand, given that the 2.0-Å radius at -2’ is slightly larger than that of a dehydrated chloride ion or desensitized canonical receptors, we cannot rule out that our GABA-bound structure represents a relatively constricted but marginally conductive state, particularly given that the conductance of ρ1 in single-channel recordings is almost two orders of magnitudes lower than for canonical receptors (Everitt et al., 2004; Kim et al., 2020; Masiulis et al., 2019; Wotring et al., 1999).

Our structures further reveal subtype-specific features that rationalize the unique pharmacology of ρ1, with likely relevance for tonic and peripheral GABA signaling (Jones and Palmer, 2009). For instance, an interfacial TMD site in canonical neuronal GABA_A_Rs is occluded by the bulky residues I15’ and W349, likely accounting for ρ1 resistance to allosteric GABA_A_R modulators such as barbiturates and benzodiazepines (Amin, 1999; Belelli et al., 1999) (Figure S6D). Similarly, loop C in ρ1 is extended by one amino acid residue, and associated with a more compact orthosteric pocket. Relative occlusion of this pocket in the ρ1 resting state, both by loop C and steric influence of Y127, prefers binding of TPMPA over the larger antagonist bicuculline (Figure 6B, top). This relative compaction appears to limit GABA access to the orthosteric site, but also to facilitate its stable coordination once bound (Figure 6B, bottom), consistent with its slow but high-affinity activation (Chang and Weiss, 1999; Polenzani et al., 1991).

Modest local rearrangements at the orthosteric site upon GABA binding propagate to relatively large domain-level transitions, of similar or greater magnitude compared to canonical neuronal subtypes—including a substantially wider outer pore (above 2’, Figure S5E). In contrast to at least some canonical GABA_A_Rs (Kim et al., 2020; Masiulis et al., 2019), the expanded activation gate in ρ1 appears to provide limited stabilizing contacts for the pore blocker PTX (Figure S7C). Accordingly, PTX preferentially stabilizes a resting-state pore, to the point of uncoupling gating transitions in the ECD from those in the TMD. This model is consistent with the mixed competitive and non-competitive effects of PTX in ρ1 functional experiments (Figure S7E). Interestingly, this mixed inhibitory profile distinguishes ρ1 even from the closely related ρ2 GABA_A_R, offering a basis for highly subtype-specific pharmaceutical targeting.

Taken together, this work elucidates structural and dynamic features of ρ1 accounting for differences in both function and pharmacology compared to other GABA_A_R subtypes. In addition to expanding our understanding of common and divergent mechanisms in this critical receptor family, these models offer templates for structure-based drug design to better target ρ1, a subtype resistant to nearly all classical drugs targeting GABA_A_Rs.

## Author Contributions

Conceptualisation: JC, CF, RJH; methodology: JC, CF, NH; investigation: JC, CF, NH, VT, YZ; data curation: CF, NH, VT; writing - original draft: JC, CF, NH; writing - review and editing: JC, CF, NH, VT, YZ, RJH, EL; funding acquisition: JC, NH, EL; supervision: RJH, EL.

## Data Availability

Cryo-EM density maps have been deposited in the Electron Microscopy Data Bank under accession numbers EMD-17106 (apo), EMD-17107 (TPMPA), EMD-17108 (PTX), EMD-17110 (GABA & PTX), and EMD-17045 (GABA). Model coordinates have been deposited in the Protein Data Bank under accession numbers PDB-8OQ6 (apo), PDB-8OQ7 (TPMPA), PDB-8OQ8 (PTX), PDB-8OQA (GABA & PTX), and PDB-8OP9 (GABA). MD simulation trajectories were deposited into Zenodo (https://doi.org/10.5281/zenodo.7807818).

## Acknowledgements

We thank Yongchang Chang, Ryan Hibbs, Colleen Noviello, and members of Molecular Biophysics Stockholm for feedback on the project and manuscript, and staff at the Swedish National Cryo-EM Facility for data collection and support. The data was collected at the Cryo-EM Swedish National Facility funded by the Knut and Alice Wallenberg, Family Erling Persson and Kempe Foundations, SciLifeLab and Stockholm University. MD simulations were performed using the computing facilities of Karolina and LUMI Supercomputer through EuroHPC (grant no. EHPC-REG-2021R0074 and EHPC-REG-2022R03-223, respectively) and Swedish National Infrastructure for Computing (SNIC 2022/3-40), and supported by BioExcel (EuroHPC grant no. 101093290). JC was supported by an EMBO Postdoctoral Fellowship, CF by grant FV-5.1.2-0523-19 from Stockholm University, NH by a Marie Sklodowska-Curie Postdoctoral Fellowship, and YZ, RJH, and EL by grants from the Swedish Research Council (2019-02433, 2021-05806) and Swedish e-Science Research Center.

## STAR METHODS

### Construct design for cryo-EM

The full-length sequence of human ρ1 GABA_A_R (uniprot: P24046) was modelled with ColabFold v1.2.0. (Jumper et al., 2021; Mirdita et al., 2022) for prediction of homopentameric structure. The number of recycles was set to 3; MMseqs2 (Mirdita et al., 2019) was used for fast homology search. Five structures were generated, relaxed (Eastman et al., 2017), and ranked based on predicted local distance difference test score. All five structures contained regions of low confidence at the N-terminus and in the predicted ICD. Based on these predictions, we designed a ρ1-EM GABA_A_R construct containing residues S58–L383 and D451–S479, separated by a sequence encoding the superfolder green fluorescent protein (Pédelacq et al., 2006). Prior to the N-terminal truncation site, we added a membrane-translocation signal peptide from the human β3 GABA_A_R, a Twin-Strep tag, and a recognition site for tobacco etch virus for expression and purification purposes (Schmidt et al., 2013).

### Expression in oocytes and electrophysiology

Plasmids encoding ρ1-WT or ρ1-EM GABA_A_R constructs were linearized using NotI or KpnI respectively, and RNA was produced by in vitro transcription using a mMessage mMachine T7 Ultra transcription kit (Ambion) according to the manufacturer protocol. The S264 mutation was introduced into the ρ1-WT background using QuikChange II mutagenesis (Agilent) and verified by sequencing. Stage IV oocytes from Xenopus laevis frogs (Ecocyte Bioscience) were injected with 12-40 ng RNA and incubated for 3–4 days at 13°C in post-injection solution (88 mM NaCl, 10 mM HEPES, 2.4 mM NaHCO_3_, 1 mM KCl, 0.91 mM CaCl_2_, 0.82 mM MgSO_4_, 0.33 mM Ca(NO_3_)_2_, 2 mM sodium pyruvate, 0.5 mM theophylline, 0.1 mM gentamicin, 17 mM streptomycin, 10,000 u/L penicillin, pH 8.5) prior to use in two-electrode voltage clamp (TEVC) measurements.

For TEVC recordings, glass electrodes were pulled and filled with 3 M KCl to give a resistance of 0.5–1.5 MΩ and used to clamp the membrane potential of injected oocytes at −80 mV with an OC-725C voltage clamp (Warner Instruments). Oocytes were maintained under continuous perfusion with Ringer’s solution (123 mM NaCl, 10 mM HEPES, 2 mM KCl, 2 mM MgSO_4_, 2 mM CaCl_2_, pH 7.5) at a flow rate of 1.5 mL/min. For dose-response curves, buffer exchange was performed by manually switching the inlet of the perfusion system into the appropriate buffer. For kinetic analysis, buffer exchange was performed using a gravity-fed, digitally-controlled solution exchange system (Scientific Instruments). Currents were digitized at a sampling rate of 2 kHz and lowpass filtered at 10 Hz with an Axon CNS 1440A Digidata system controlled by pCLAMP 10 (Molecular Devices).

For dose-response curves, current responses were measured at the end of a 2-min pulse of agonist, or at the maximal current amplitude during the pulse for traces showing desensitization. Dose-response curves were fitted by nonlinear regression to the Boltzmann equation with variable slope and amplitudes using Prism 9.4 (GraphPad Software). Inhibition by TPMPA and bicuculline was measured by 100-second pulses of the respective compound during a 300-second pulse of 1 μM GABA. Inhibition was reported as the current remaining at the end of the 100-second pulse as a fraction of the average current at the end of the flanking 100-second exposures to 1 μM GABA alone. Each reported value represents the mean and the standard error of the mean for three to four oocytes.

### Baculovirus production

Chemically competent DH10BacVSV cells (Geneva Biotech) were transformed with a modified pEZT-BM vector encoding the ρ1-EM GABA_A_R and incubated at 37° C on transposition plates until the blue-white distinction was apparent (Morales-Perez et al., 2016). White colonies were picked and grown in suspension overnight and bacmid DNA was isolated as described previously (Goehring et al., 2014). Bacmids encoding ρ1-EM GABA_A_Rs were used to transfect TriEX Sf9 cells (Novagen) in 6-well plates using ExpiFectamine-SF (Fisher) according to manufacturer instructions. Supernatant containing the P1 viruses was harvested following 96 h incubation. Virus was amplified by infecting a 30-mL suspension Sf9 cell culture as described previously to generate P2 virus (Goehring et al., 2014). P2 virus was further amplified to 300 mL by infecting large cultures of Sf9 cells to generate P3 virus that was used for infecting HEK cells.

### Expression and purification

Baculovirus encoding the ρ1-EM GABA_A_R was used to infect Expi293F GnTl^-^ cells for protein production at 2.5% v/v virus-to-cell culture at a density of 2 × 10^6^ cells/mL. After 6 h incubation at 37°C, 3 mM valproic acid was added to cultures and flasks were moved to 30°C. Cells were harvested 48 h post-transduction and washed with phosphate buffered saline prior to flash-freezing cell pellets.

All steps of protein purification were performed at 4°C. Cell pellets (∼40g wet mass) from 2 L culture were resuspended in 100 mL resuspension buffer (300 mM NaCl, 40 mM HEPES pH 7.5 and 2 cOmplete protease inhibitor tablets (Roche)) and sonicated to disrupt cell membranes. Lysed cells were spun at 50,000 x g for 45 min to pellet membranes. The membrane pellet was resuspended in 100 mL 2X solubilization buffer (600 mM NaCl, 80 mM HEPES, pH 7.5 with 2 tablets of cOmplete protease inhibitor) and homogenized using a Dounce homogenizer. Solubilization was initiated by addition of 100 mL 2X detergent mixture (2% lauryl maltose neopentyl glycol (LMNG), 0.2% cholesteryl hemisuccinate (CHS)) and gentle stirring for 2 h and 40 min. Insoluble debris was pelleted by centrifugation at 50,000 x g for 55 min. The supernatant was applied to 4 mL Streptactin Superflow resin (IBA) that was previously equilibrated in Buffer A (300 mM NaCl, 20 mM HEPES, 0.005% LMNG, 0.0005% CHS, pH 7.5) and batch-bound with gentile mixing for 90 min. Resin was washed with 30 column volumes of Buffer A and eluted using 4 column volumes Buffer A with 10 mM desthiobiotin (Sigma). The affinity-purified protein was concentrated and further purified by gel-filtration chromatography on a Superose 6 increase 10/300 column (Cytiva) with a mobile phase of 100 mM NaCl, 20 mM HEPES, 0.005% LMNG, 0.0005% CHS, pH 7.5 with or without 50 μM GABA. Peak fractions were pooled and concentrated for nanodisc reconstitution.

### Nanodisc reconstitution

The plasmid for SapA expression was a gift from Salipro Biotech AB. Purification of SapA followed previously published protocols (Lyons et al., 2017). For the reconstitution of saposin nanodiscs, purified ρ1-EM GABA_A_Rs, SapA and polar brain lipid (Avanti) were mixed as molar ratio 1:15:150, then incubated on ice for 1 h. After that, Bio-Beads SM-2 resin (Bio-Rad) was added and the mixture was gently shaken overnight at 4°C. On the next day, the supernatant containing SapA nanodiscs was collected and further purified by gel-filtration chromatography on a Superose 6 column with buffer containing 20 mM HEPES pH 7.5, 100 mM NaCl. Peak fractions were pooled and concentrated to ∼3.5 mg/mL.

### Grid preparation and EM data acquisition

In order to minimize preferred orientation during freezing, fluorinated foscholine 8 (FFC-8) (Anatrace) was added to nanodiscs before grid freezing. A concentrated nanodisc solution at ∼3.5 mg/mL (2.7 μL) was combined with 10X stock of FFC-8 (0.3 μL) and ligands, to give final concentrations of 3 mM FFC-8 (apo dataset), 3 mM FFC-8 with 1 mM TPMPA (TPMPA dataset), 2 mM FFC-8 with 100 μM PTX (PTX dataset), 3 mM FFC-8 with 200 μM GABA (GABA dataset), or 2 mM FFC-8 with 100 μM PTX and 400 μM GABA (GABA & PTX dataset). For each grid, 3 μL of the mixture was applied to a glow-discharged R1.2/1.3 400 mesh Au grid (Quantifoil), blotted for 2 s with force 0 and plunged into liquid ethane using a Vitrobot Mark IV (Thermo Fisher Scientific) at 20°C. Cryo-EM data were collected at 300 kV using a Titan Krios (Thermo Fisher Scientific) electron microscope fitted with a K3 Summit detector (Gatan) operating at a 15 e^-^/px/s flux and a magnification corresponding to 0.6645 or 0.6725 Å/px using the EPU automated collection software (Thermo Fisher Scientific). Micrographs were recorded over a period of 1.2 s and subdivided into 40 frames for a total fluence of ∼45e^-^/Å^2^ at the sample level.

### Image processing

Dose-fractionated images in super-resolution mode were internally gain-normalized and binned by 2 in EPU during data collection. Motion correction of the dose-fractionated images was performed in Relion 4.0 (Kimanius et al., 2021). The contrast transfer functions (CTF) were estimated for motion-corrected images using CTFFIND4.1 (Grant et al., 2018) and particles were automatically picked using Topaz 0.2.5 by the internal trained model (Bepler et al., 2019). After two rounds of 2D classification in Relion 4.0 to remove junk particles, good classes were picked, re-extracted and centered by applying alignment offsets. These particles were used to generate initial models in CryoSPARC 4.0 (Punjani et al., 2020). Further processing was done in Relion 4.0, including 3D classification to check the structural heterogeneity of classes and further clean the dataset. Multiple rounds of CtfRefine and polishing were executed to improve resolution. The apo, TPMPA, and GABA structures were refined with C5 symmetry, while the PTX and GABA & PTX structures were refined without symmetry imposed.

### Model building

Initial models of ρ1-EM GABA_A_R monomers were created using AlphaFold2 as described above, fitted using Coot (Emsley et al., 2010) and manually edited to improve the local fit to density. Crystallographic information files (cif) for ligands were generated from isomeric SMILES strings using Grade2 (Smart O.S. et al., 2021). Initial monomer models were refined using Rosetta fast_relax (Khatib et al., 2011) with “electron_density_fast” score reweighting (DiMaio et al., 2009). Ligand parameter files were generated from cifs using conformers calculated in Open Babel3 (Yoshikawa and Hutchison, 2019) and converted using the molfile_to_params.py script in Rosetta. Glycans were modeled as a single basal N-acetyl glucosamine moiety and connected using link records. Where relevant, symmetry was generated by manual fitting of a neighbor monomer copy, and creation of a symmetry definition file using make_symmdef_file_denovo.py in Rosetta. Symmetrized models were then subjected to a second round of Rosetta fast_relax.

Final models were optimized using real-space refinement in PHENIX (Adams et al., 2010) and validated by MolProbity (Williams et al., 2018). Side chains were truncated for residues without clear densities, mostly in the ICD. Subunit interfaces were analyzed using the PDBePISA server (Krissinel and Henrick, 2007), and pore radius profiles were calculated using HOLE (Smart et al., 1996). Structure figures were prepared using UCSF ChimeraX (Pettersen et al., 2021) and Pymol (DeLano, 2002).

### Molecular dynamics simulations

Simulations were performed by embedding each protein system in a symmetric membrane that approximates neuronal plasma membrane composition: 44.4% cholesterol, 22.2% 1-palmitoyl-2-oleoyl-sn-glycero-3-phosphocholine (POPC), 22.2% 1-palmitoyl-2-oleoyl-sn-glycero-3-phosphoethanolamine (POPE), 10% 1-palmitoyl-2-oleoyl-sn-glycero-3-phospho-L-serine (POPS) and 1.1% phosphatidylinositol 4,5-bisphosphate (PtdIns(4,5)P_2_) (Ingólfsson et al., 2017). The systems were built using the Membrane Builder module of CHARMM-GUI (Jo et al., 2008). Each system was solvated with TIP3P water (Jorgensen et al., 1983) and neutralized in 0.15 M KCl to generate systems containing ∼300, 000 atoms each, with dimensions of 130 × 130 × 170 Å^3^. To allow for better sampling, six independent replicas of each system were built by randomly configuring initial lipid placement around the protein-ligand system (Licari et al., 2022). To obtain comparable sampling of ligand-protein interactions in the canonical neuronal receptor system (PDB ID 6X3Z), which contains only two GABA binding sites compared to five in ρ1, six more replicates were generated. Following the default CHARMM-GUI settings, each system was energy minimized and then relaxed in simulations at constant pressure (1 bar) and temperature (300K) for 30 ns, during which the position restraints on the protein were gradually released.

MD simulations in this study were performed using GROMACS-2023 (Páll et al., 2020) utilizing CHARMM36m (Huang et al., 2017) force field parameters for proteins and lipids, respectively. The force field parameters for all the ligands were generated using the CHARMM General Force Field (CGenFF) (Vanommeslaeghe et al., 2012, 2010; Vanommeslaeghe and MacKerell, 2012). Cation-π interaction specific NBFIX parameters were used to maintain appropriate ligand-protein interactions at the aromatic cage, located at the orthosteric binding site (Liu et al., 2021). Bonded and short-range nonbonded interactions were calculated every 2 fs, and periodic boundary conditions were employed in all three dimensions. The particle mesh Ewald (PME) method (Darden et al., 1993) was used to calculate long-range electrostatic interactions with a grid density of 0.1 nm^−3^. A force-based smoothing function was employed for pairwise nonbonded interactions at 1 nm with a cutoff of 1.2 nm. Pairs of atoms whose interactions were evaluated were searched and updated every 20 steps. A cutoff (1.2 nm) slightly longer than the nonbonded cutoff was applied to search for the interacting atom pairs. Constant pressure was maintained at 1 bar using the Parrinello-Rahman algorithm (Parrinello and Rahman, 1980). Temperature coupling was kept at 300K with the v-rescale algorithm (Bussi et al., 2007).

### Analysis

Upper-ECD spread, β-expansion, and domain twist were calculated using the definitions introduced by Lev and colleagues (Lev et al.., 2017). Briefly, upper-ECD spread was defined as the distance from center-of-mass (COM) of residues 74 to 104, 122 to 160, 168 to 192, 216 to 237, and 258 to 271 of each subunit to the same of the entire pentamer. β-expansion was defined as the distance between the COM of residues 111 to 115 to that of residues 277 to 281. Domain twist was the dihedral angle formed by four positions: the COM of the ECD of a subunit, COM of the ECD of the pentamer, COM of the TMD of the pentamer and COM of the TMD of a submit.

## SUPPLEMENTAL INFORMATION

**Figure S1.**
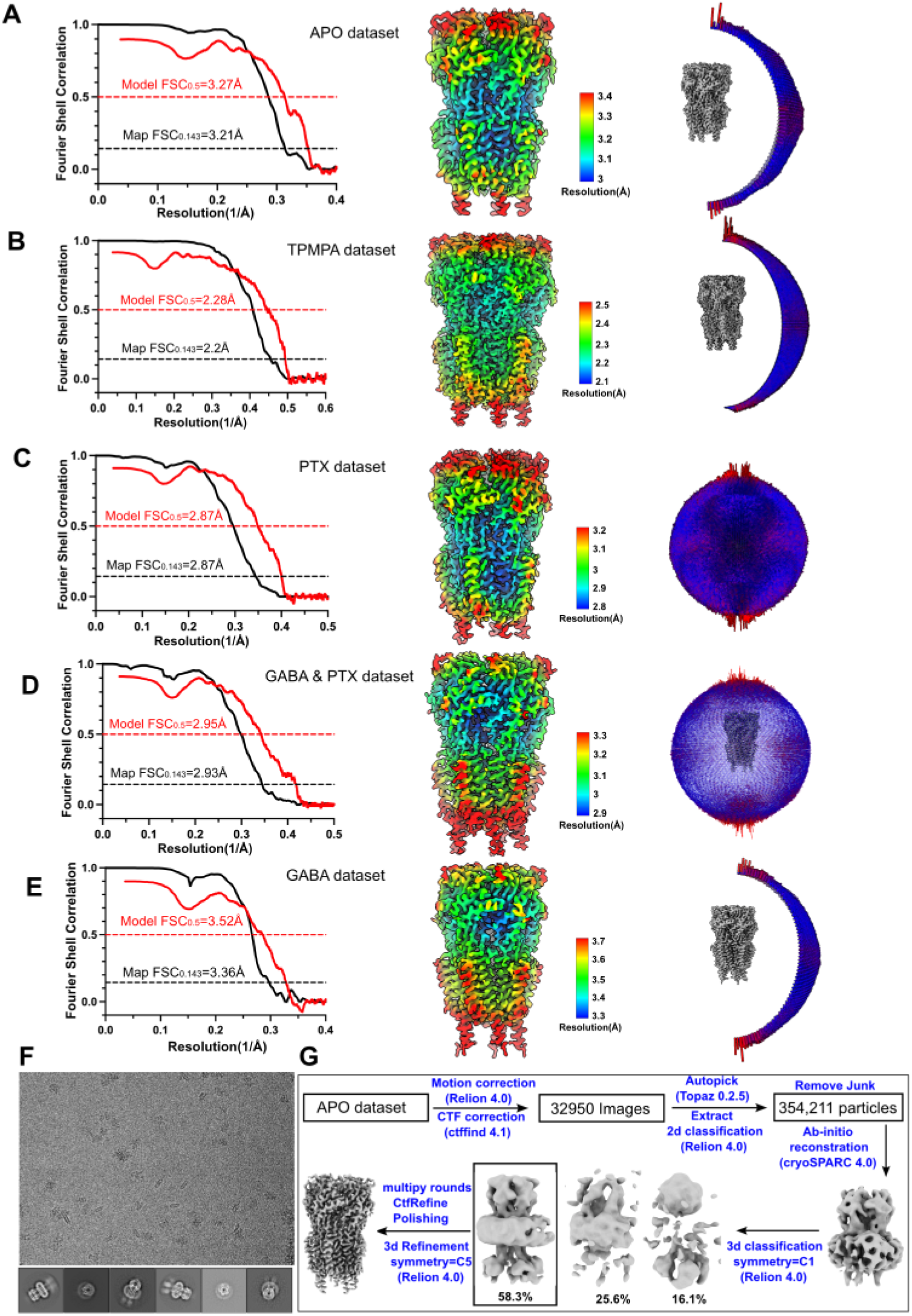
Processing pipelines for ρ1-EM GABA_A_R structures. (A–E) Processing statistics, local resolution, and angular distributions in cryo-EM data producing structures of ρ1-EM GABA_A_Rs under apo (A), TPMPA (B), PTX (C), PTX & GABA (D), or GABA alone (E) conditions. Each panel shows *left:* Fourier-shell correlations (FSCs) between two independently refined half-maps after masking (black) and from cross validation of the atomic model against the final cryo-EM map (red); *center:* final cryo-EM map coloured by local resolution; *right:* angular distribution of particles used in the cryo-EM reconstitution. (F) Representative cryo-EM image and 2D classification images from the apo dataset. (G) Example cryo-EM data processing workflow for the apo dataset.

**Figure S2.**
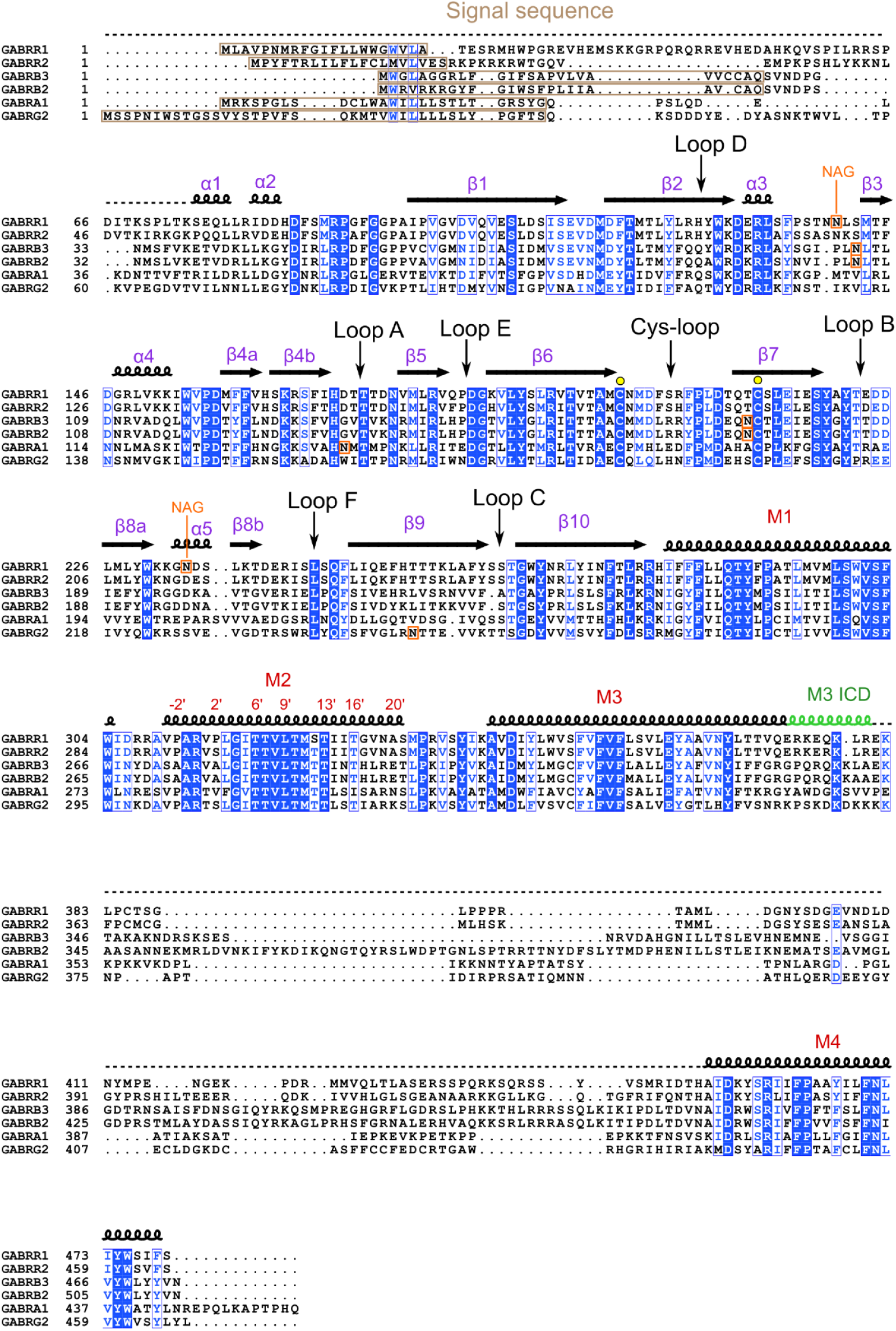
Alignment of amino acid sequences of human ρ versus canonical neuronal GABA_A_R subtypes. Sequence alignment of human ρ1, ρ2, β3, β2, α1 and γ2 GABA_A_R subtypes. Residues are numbered according to reference UniProt sequences. Key structural features are labeled above alignment, including the signal sequence (brown), glycosylation sites (orange), disulfide bridge (yellow), secondary structure elements of the ECD (purple) and TMD (red), and the resolved region of the ICD (green). Blue highlights indicate residues that are fully (filled) or partially (open) conserved among the selected subtypes.

**Figure S3.**
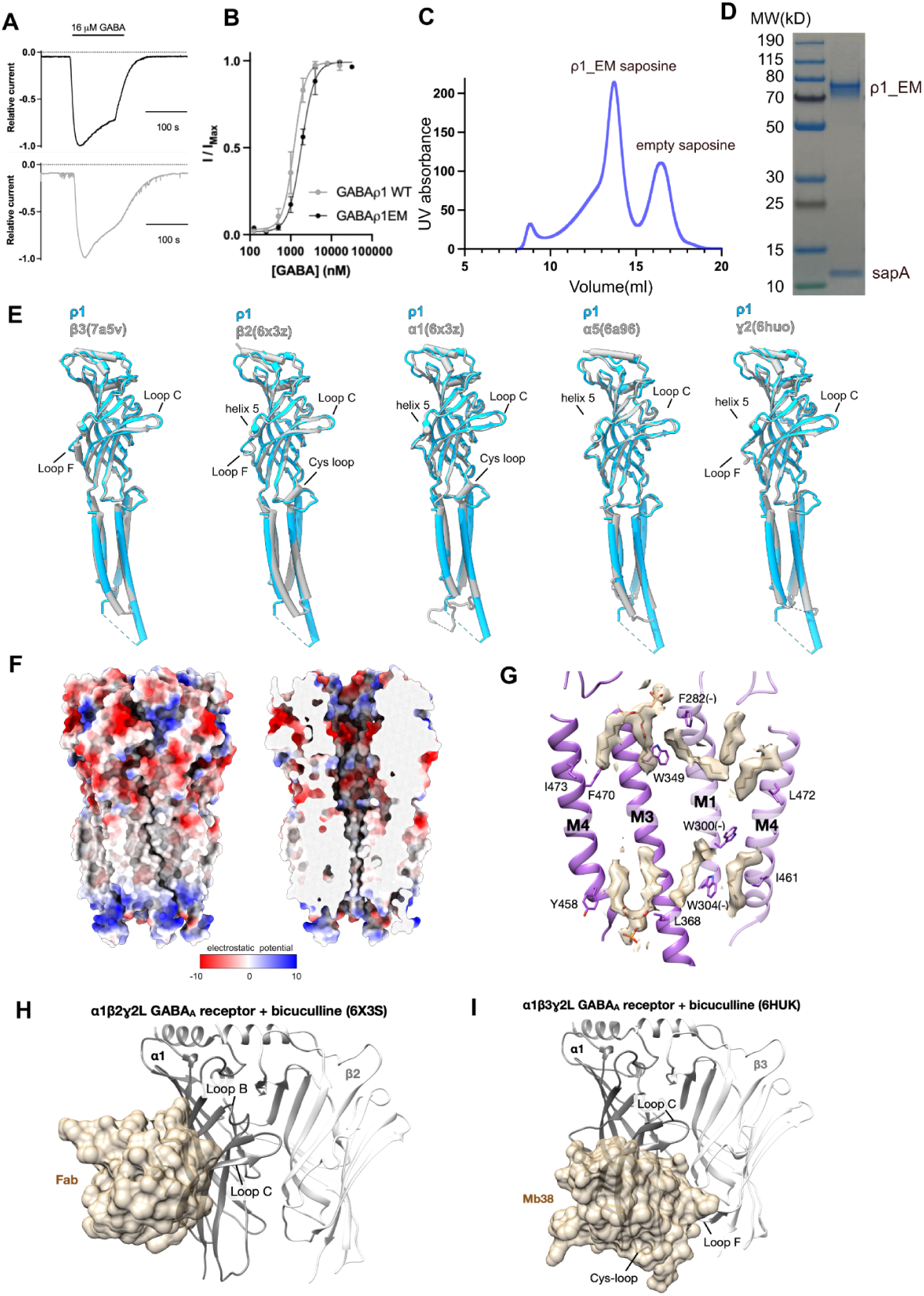
Function, biochemical and structural features of ρ1-EM and related GABA_A_Rs. (A) Example traces from TEVC recordings of p1-EM (top, black) and -WT (bottom, gray) GABA_A_Rs in response to 16-μM GABA applications for 100 seconds in *X. laevis* oocytes. (B) Concentration-dependence curves for p1-EM (black) and -WT GABA_A_Rs in response to pulses of GABA up to 2 min. Bars represent standard errors of the mean (SEM) from measurements of 4 (p1-EM) or 5 (p1-WT) oocytes. (C) Size-exclusion chromatography (SEC) of ρ1-EM GABA_A_Rs reconstituted in saposin-brain lipid nanodiscs. (D) Gel electrophoresis (SDS-PAGE) of ρ1-EM GABA_A_Rs reconstituted in saposin nanodiscs. Bands corresponding to the predicted sizes of GABA_A_R complexes (ρ1-EM) and empty nanodiscs (sapA) are labeled accordingly. (E) Overlay of a single GABA-bound p1-EM GABA_A_R subunit (blue) with individual subunits from structures of other GABA_A_R subtypes solved previously by cryo-EM (gray). Comparative isoforms are *left–right:* β3 (from β3 homopentamer with histamine & Mb25, PDB ID 7A5V), β2 or α1 (from α1β2ɣ2 with GABA & 1F4-Fab, PDB ID 6X3Z), α5 (from α5β3 with GABA & Nb25, PDB ID 6A96), or ɣ2 (from α1β3ɣ2L with GABA, alprazolam & Mb38, PDB ID 6HUO). (F) Surface representations of the apo structure of ρ1, colored by electrostatic potential, showing *left:* full receptor or *right:* cutaway through the pore axis. (G) Overlay of non-protein cryo-EM densities (brown) around lipid molecules built in the apo structure of ρ1, with nearby residues of the principal (dark purple) and complementary (light purple) subunits shown as sticks and labeled accordingly. (H–I) Fiduciary labels bound to inhibited states of α1β2γ2 (H) and α1β3γ2L (I) structures. The Fab fragment or megabody is shown as a semi-transparent surface (brown), with the receptor model shown as ribbons (gray). For clarity, only ECDs from two subunits are shown.

**Figure S4.**
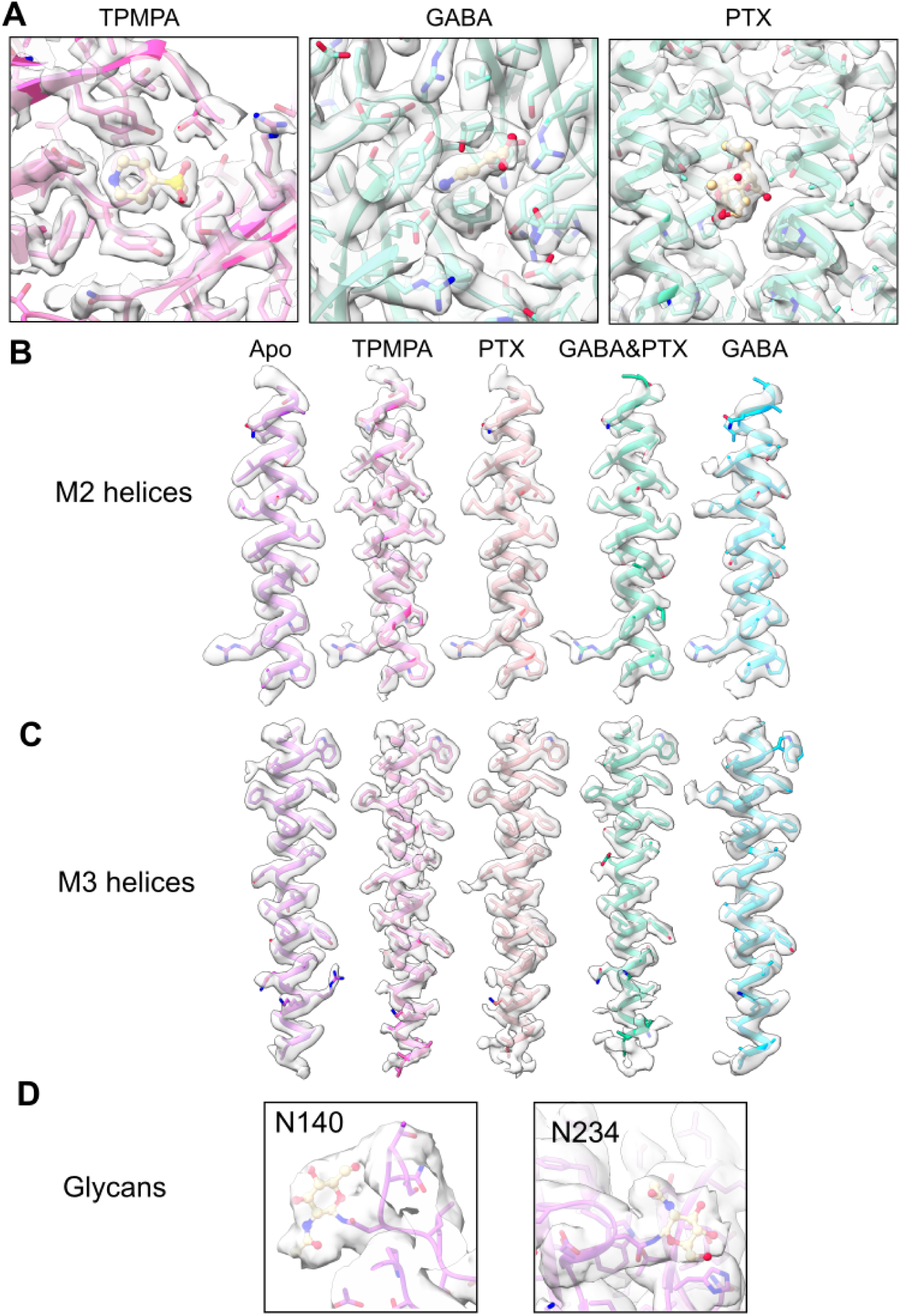
Representative densities around key ligand, protein, and glycosylation sites in ρ1-EM GABA_A_Rs. (A) Densities and models of the ligand and surrounding protein for *left:* TPMPA, *center:* GABA (middle) or *right:* PTX. (B) Densities and models of M2 helices in all ρ1-EM GABA_A_R structures in this study. (C) Densities and models of M3 helices in all ρ1-EM GABA_A_R structures in this study. (D) Densities and models of glycans in the apo structure of the ρ1-EM GABA_A_R.

**Figure S5.**
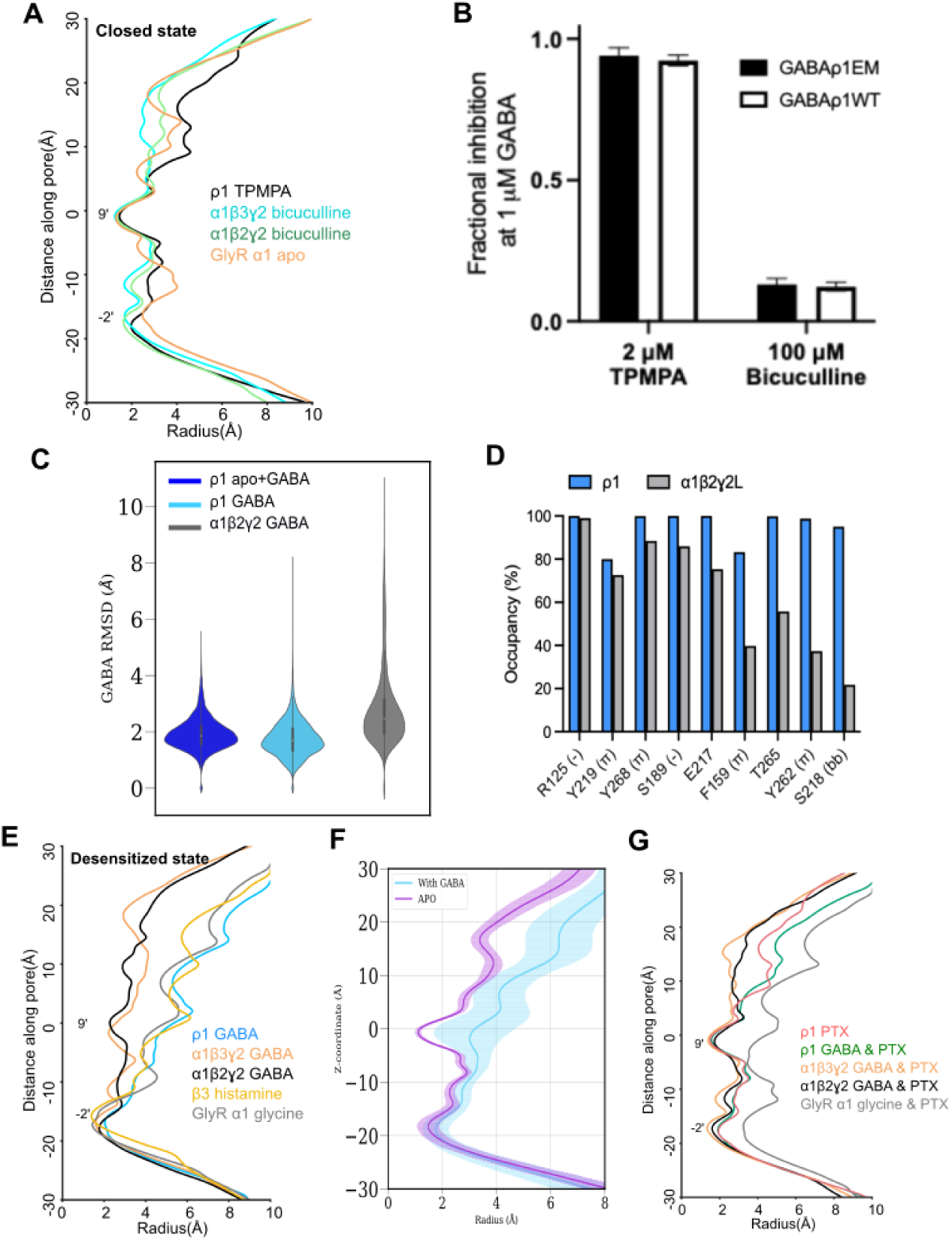
Comparative pore profiles, drug inhibition, and ligand dynamics of ρ1 versus related GABA_A_Rs. (A) Pore-radius profiles from representative GABA_A_R and GlyR structures in presumed resting states, including ρ1 with TPMPA (black), α1β3γ2L GABA_A_R with bicuculline & Mb38 (PDB ID 6HUK, cyan), α1β2γ2 GABA_A_R with bicuculline & 1F4-Fab (PDB ID 6X3S, green), and α1 GlyR (PDB ID 6UBS, orange). (B) Fractional inhibition of ρ1-EM (black) and -WT (white) GABA_A_R currents in the presence of 1 μM GABA by TPMPA (2 μM) or bicuculline (100 μM). Fractional inhibition is the fraction of the maximal (baseline-subtracted) current amplitude during treatment with GABA alone that remains at the end of the treatment with TPMPA or bicuculline. Bars represent SEM from 4 independent oocytes for each experiment. (C) Violin plots of GABA RMSD (Å) during all MD-simulation replicates, calculated by first aligning the ECD of each subunit to the starting structure. (D) Percent occupancies of the nine most prevalent hydrogen-bond or cation-π interactions of GABA with protein residues in MD simulations of GABA-bound ρ1 (blue) or α1β2ɣ2 (PDB ID 6X3Z, gray) GABA_A_Rs. A hydrogen bond was counted to be formed by a hydrogen atom (H) covalently bound to an electronegative donor atom (D), interacting with an electronegative acceptor atom (A), provided that the distance D–A is <3 Å and the angle D–H–A is more than 120°. A cation-π interaction was counted to be formed if the distance between the GABA nitrogen atom and the center-of-mass of the aromatic ring of a protein residue is <6 Å. (E) Pore-radius profiles from representative GABA_A_R and GlyR structures in presumed desensitized states, including ρ1 with GABA (blue), α1β3γ2L GABA_A_R with GABA, diazepam & Mb38 (PDB ID 6HUP, orange), α1β2γ2 GABA_A_R with GABA & 1F4-Fab (PDB ID 6X3Z, black), β3 GABA_A_R with histamine & Mb25 (PDB ID 7A5V, yellow), and α1 GlyR with glycine (PDB ID 6PLR, gray). (F) Pore-radius profiles during MD simulations of ρ1 in resting (purple) and desensitized (cyan) states, calculated using the software package CHAP. Solid lines represent simulation means, shaded regions indicate standard deviations. (H) Pore-radius profiles from representative GABA_A_R and GlyR structures bound to PTX, including ρ1 with PTX (red), ρ1 with GABA & PTX (green), α1β3γ2L GABA_A_R with GABA, PTX & Mb38 (PDB ID 6HUJ, orange), α1β2γ2 GABA_A_R with GABA, PTX & 1F4-Fab (PDB ID 6X40, black), and α1 GlyR with glycine & PTX (PDB ID 6UD3, gray).

**Figure S6.**
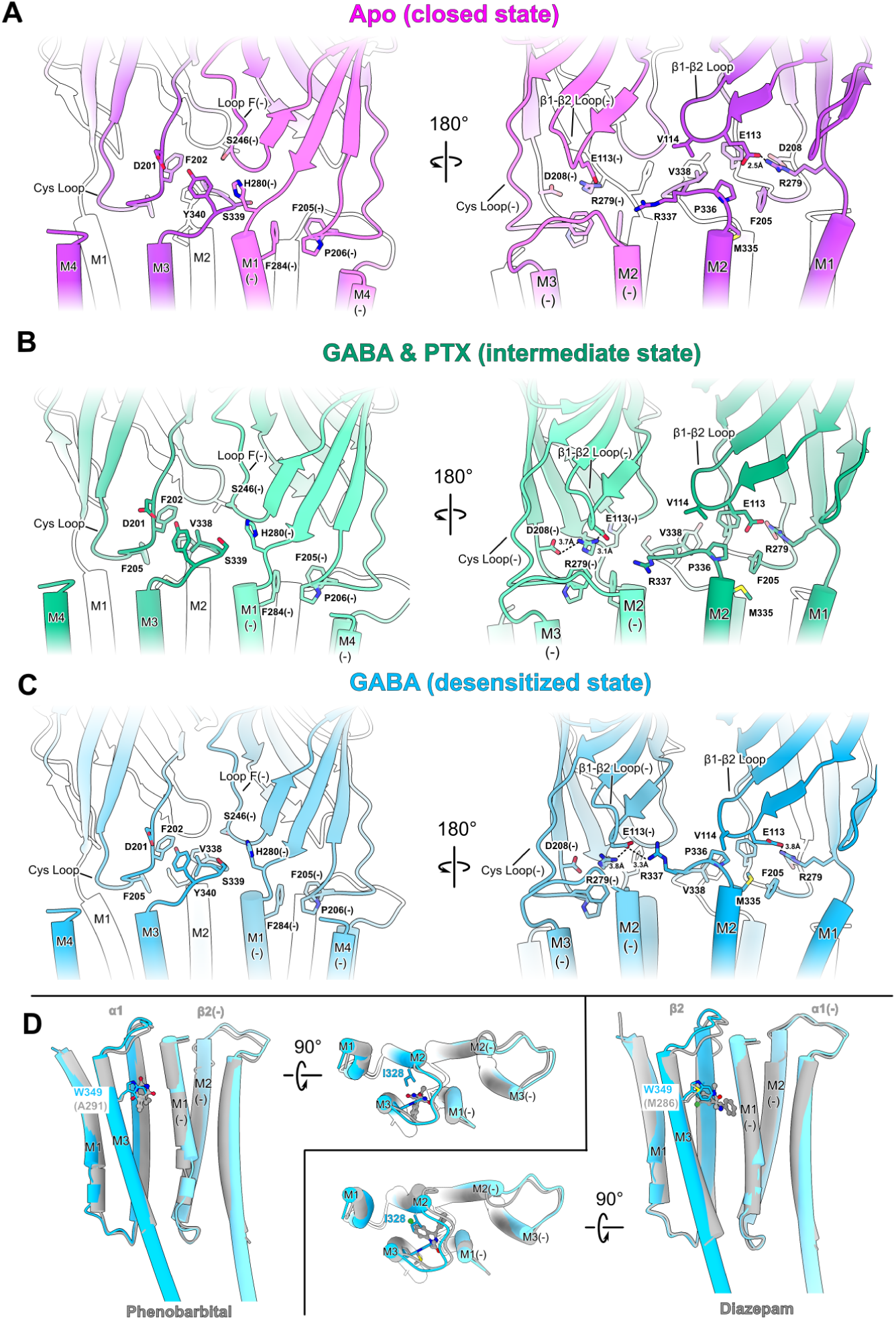
Amino acid contacts at domain and subunit interfaces. (A–C) Zoom views of the ECD-TMD interface of ρ1 in resting (TPMPA structure, pink), intermediate (GABA & PTX structure, green), and desensitized (GABA structure, cyan) states. For clarity, each view shows the interface between a single principal (dark) and complementary (light) subunit. Key amino-acid residues are labeled and shown as sticks. (D) Superposition of TMD drug-binding sites in α1β2ɣ2 GABA_A_Rs (gray) with GABA-bound ρ1 (cyan), viewed from either the membrane plane or extracellular side. Comparative structures include α1β2ɣ2 GABA_A_Rs in the presence of GABA & 1F4-Fab with *left:* phenobarbital (αβ interface, PDB ID 6X3W) or *right:* diazepam (βα interface, PDB ID 6X3W). Key residues M2-I328 and M3-W349 on the principal ρ1 subunit are labeled and shown as cyan sticks; drugs are shown as gray balls-and-sticks, colored by heteroatom.

**Figure S7.**
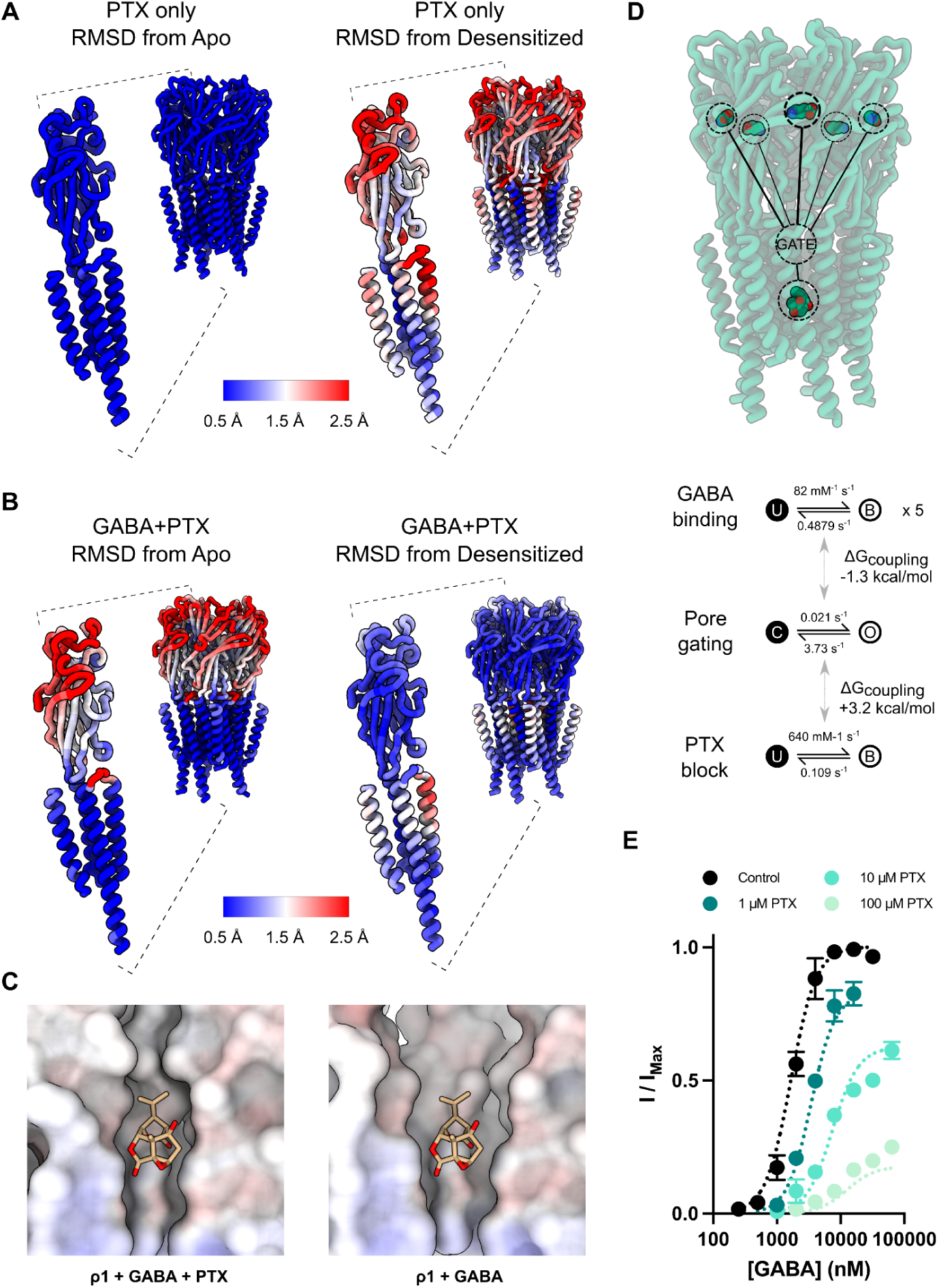
PTX uncoupling of GABA-induced pore expansion. (A–B) Structures of ρ1 with PTX (A) or GABA & PTX (B), viewed from the membrane plane and colored according to the central scale bar by RMSD relative to the *left:* resting or *right:* desensitized states, based on alignment on the M1 helix (residues 285–299). *Inset:* zoom view of a single subunit. (C) Models of PTX (brown sticks, colored by heteroatom) in the *left:* resting or *right:* desensitized ρ1 pore (surface, colored by electrostatic potential). For clarity, the proximal two protein subunits are not shown. PTX in the resting pore represents the PTX structure; PTX in the desensitized state was manually inserted based on alignment of the M2 helices in the PTX structure with those in the GABA structure. (D) Structural (top) and elemental (bottom) depiction of an allosteric model of GABA and PTX action on channel gating, assuming 5 identical GABA-binding sites. In this model, PTX can bind to and block either the closed or open pore, but preferentially stabilizes the closed pore, as shown by the positive coupling energy term between the PTX element and pore gate. (E) Experimental (circles) and simulated (dotted lines) GABA concentration-response curves at varying concentrations of PTX, showing both a rightward shift in apparent affinity and a reduction in efficacy of GABA at increasing concentrations of PTX, characteristic of an allosteric inhibitor. Bars represent SEM from 3 independent oocytes.

**Video S1 | Representative states of the ρ1 GABAAR**

Morph from the apo structure to the GABA & PTX intermediate, then to the structure with GABA alone, then back to the apo structure. Transitions are first shown from the extracellular side, then rotated to show views from the membrane plane. A single subunit is highlighted as opaque, similar to the depiction in Figure 1.

**Video S2 | Resting and GABA-bound ρ1 and α1β2ɣ2 GABA_A_Rs**

A 360° view of apo ρ1 GABA_A_R from the membrane plane, overlaid with the bicuculline-bound α1β2ɣ2 GABA_A_R (PDB ID 6X3S) with bicuculline shown as spheres. The bicuculline structure is then replaced with the GABA-bound α1β2ɣ2 GABA_A_R (PDB ID 6X3Z), and the apo ρ1 is morphed to the GABA-bound ρ1. Finally, the GABA-bound ρ1 and α1β2ɣ2 GABA_A_R structures are compared with a 360° rotation. For the α1β2ɣ2 GABA_A_R, α1 is shown in gray, β2 is shown in white, and ɣ2 is shown in dark gray.

**Table 1.**
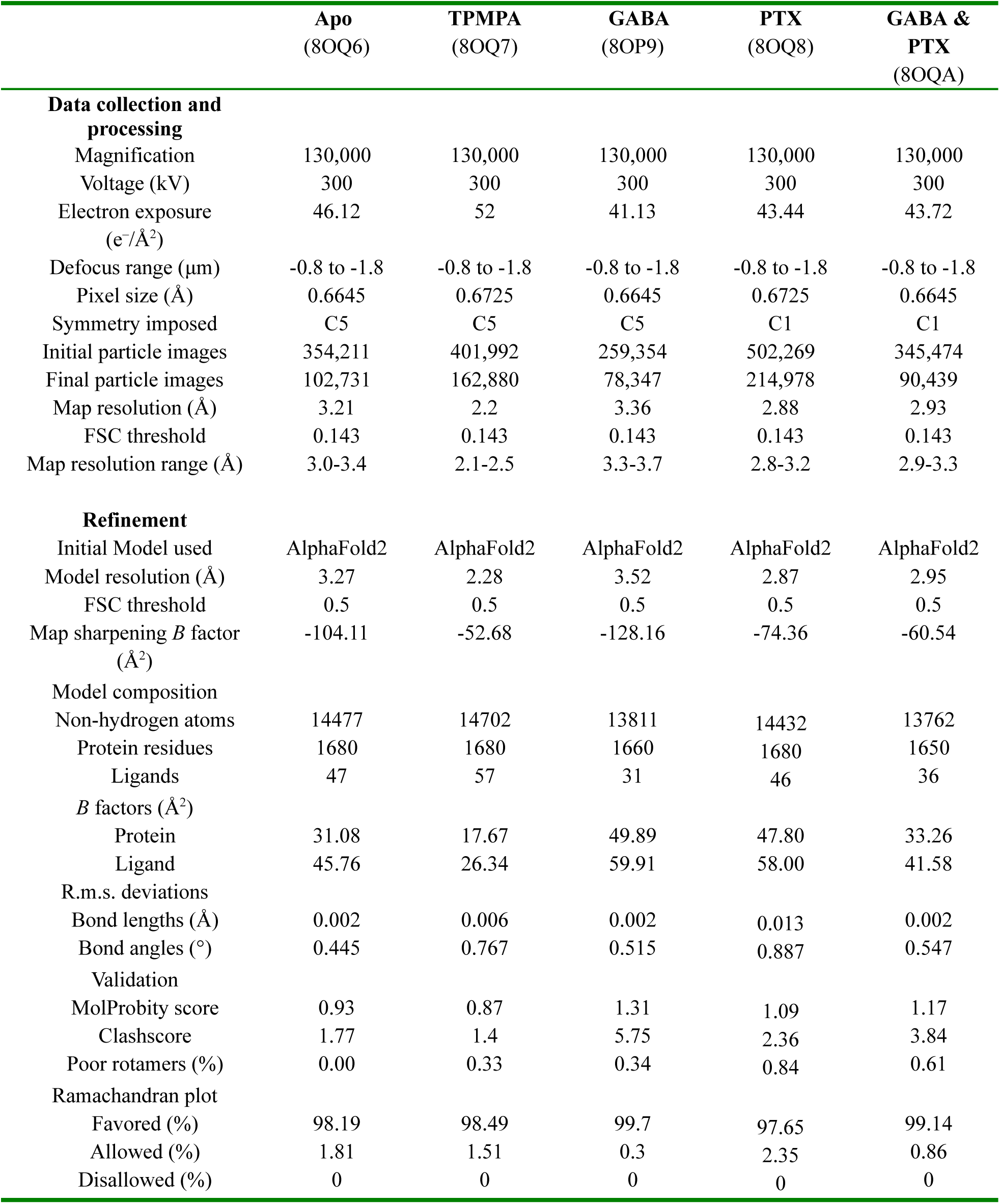
Cryo-EM data collection, refinement and validation statistics.

## REFERENCES

Adams, P.D., Afonine, P.V., Bunkóczi, G., Chen, V.B., Davis, I.W., Echols, N., Headd, J.J., Hung, L.-W., Kapral, G.J., Grosse-Kunstleve, R.W., McCoy, A.J., Moriarty, N.W., Oeffner, R., Read, R.J., Richardson, D.C., Richardson, J.S., Terwilliger, T.C., Zwart, P.H., 2010. PHENIX: a comprehensive Python-based system for macromolecular structure solution. Acta Crystallogr. D Biol. Crystallogr. 66, 213–221. https://doi.org/10.1107/S0907444909052925

Amin, J., 1999. A Single Hydrophobic Residue Confers Barbiturate Sensitivity to γ-Aminobutyric Acid Type C Receptor. Mol. Pharmacol. 55, 411.

Ananchenko, A., Hussein, T.O.K., Mody, D., Thompson, M.J., Baenziger, J.E., 2022. Recent Insight into Lipid Binding and Lipid Modulation of Pentameric Ligand-Gated Ion Channels. Biomolecules 12. https://doi.org/10.3390/biom12060814

Arnaud, C., Gauthier, P., Gottesmann, C., 2001. Study of a GABAc receptor antagonist on sleep-waking behavior in rats. Psychopharmacology (Berl.) 154, 415–419. https://doi.org/10.1007/s002130000653

Belelli, D., Pau, D., Cabras, G., Peters, J.A., Lambert, J.J., 1999. A single amino acid confers barbiturate sensitivity upon the GABA ρ1 receptor. Br. J. Pharmacol. 127, 601–604. https://doi.org/10.1038/sj.bjp.0702611

Bepler, T., Morin, A., Rapp, M., Brasch, J., Shapiro, L., Noble, A.J., Berger, B., 2019. Positive-unlabeled convolutional neural networks for particle picking in cryo-electron micrographs. Nat. Methods 16, 1153–1160. https://doi.org/10.1038/s41592-019-0575-8

Blednov, Y.A., Benavidez, J.M., Black, M., Leiter, C.R., Osterndorff-Kahanek, E., Johnson, D., Borghese, C.M., Hanrahan, J.R., Johnston, G.A.R., Chebib, M., Harris, R.A., 2014. GABAA Receptors Containing ρ1 Subunits Contribute to In Vivo Effects of Ethanol in Mice. PLOS ONE 9, e85525. https://doi.org/10.1371/journal.pone.0085525

Bormann, J., 1988. Electrophysiology of GABAA and GABAB receptor subtypes. Trends Neurosci. 11, 112–116. https://doi.org/10.1016/0166-2236(88)90156-7

Bussi, G., Donadio, D., Parrinello, M., 2007. Canonical sampling through velocity rescaling. J. Chem. Phys. 126, 014101. https://doi.org/10.1063/1.2408420

Chang, Y., Weiss, D.S., 1999. Channel opening locks agonist onto the GABAC receptor. Nat. Neurosci. 2, 219–225. https://doi.org/10.1038/6313

Colquhoun, D., 1999. GABA and the single oocyte: relating binding to gating. Nat. Neurosci. 2, 201–202. https://doi.org/10.1038/6298

Cunha, C., Monfils, M., LeDoux, J., 2010. GABAC receptors in the lateral amygdala: a possible novel target for the treatment of fear and anxiety disorders? Front. Behav. Neurosci. 4.

Cutting, G.R., Lu, L., O’Hara, B.F., Kasch, L.M., Montrose-Rafizadeh, C., Donovan, D.M., Shimada, S., Antonarakis, S.E., Guggino, W.B., Uhl, G.R., 1991. Cloning of the gamma-aminobutyric acid (GABA) rho 1 cDNA: a GABA receptor subunit highly expressed in the retina. Proc. Natl. Acad. Sci. 88, 2673–2677. https://doi.org/10.1073/pnas.88.7.2673

Darden, T., York, D., Pedersen, L., 1993. Particle mesh Ewald: An N⋅log(N) method for Ewald sums in large systems. J. Chem. Phys. 98, 10089–10092. https://doi.org/10.1063/1.464397

DeLano, W.L., 2002. The PyMOL Molecular Graphics System.

DiMaio, F., Tyka, M.D., Baker, M.L., Chiu, W., Baker, D., 2009. Refinement of protein structures into low-resolution density maps using rosetta. J. Mol. Biol. 392, 181–190. https://doi.org/10.1016/j.jmb.2009.07.008

Dong, C., Picaud, S., Werblin, F., 1994. GABA transporters and GABAC-like receptors on catfish cone-but not rod-driven horizontal cells. J. Neurosci. 14, 2648. https://doi.org/10.1523/JNEUROSCI.14-05-02648.1994

Drew, C.A., Johnston, G.A.R., Weatherby, R.P., 1984. Bicuculline-insensitive GABA receptors: Studies on the binding of (−)-baclofen to rat cerebellar membranes. Neurosci. Lett. 52, 317–321. https://doi.org/10.1016/0304-3940(84)90181-2

Eastman, P., Swails, J., Chodera, J.D., McGibbon, R.T., Zhao, Y., Beauchamp, K.A., Wang, L.-P., Simmonett, A.C., Harrigan, M.P., Stern, C.D., Wiewiora, R.P., Brooks, B.R., Pande, V.S., 2017. OpenMM 7: Rapid development of high performance algorithms for molecular dynamics. PLoS Comput. Biol. 13, e1005659. https://doi.org/10.1371/journal.pcbi.1005659

Emsley, P., Lohkamp, B., Scott, W.G., Cowtan, K., 2010. Features and development of Coot. Acta Crystallogr. D Biol. Crystallogr. 66, 486–501. https://doi.org/10.1107/S0907444910007493

Enz, R., Cutting, G.R., 1998. Molecular composition of GABAC receptors. Vision Res. 38, 1431–1441. https://doi.org/10.1016/S0042-6989(97)00277-0

Everitt, A.B., Luu, T., Cromer, B., Tierney, M.L., Birnir, B., Olsen, R.W., Gage, P.W., 2004. Conductance of Recombinant GABA Channels Is Increased in Cells Co-expressing GABAA A Receptor-associated Protein*. J. Biol. Chem. 279, 21701–21706. https://doi.org/10.1074/jbc.M312806200

Feigenspan, A., Wässle, H., Bormann, J., 1993. Pharmacology of GABA receptor CI− channels in rat retinal bipolar cells. Nature 361, 159–162. https://doi.org/10.1038/361159a0

Goehring, A., Lee, C.-H., Wang, K.H., Michel, J.C., Claxton, D.P., Baconguis, I., Althoff, T., Fischer, S., Garcia, K.C., Gouaux, E., 2014. Screening and large-scale expression of membrane proteins in mammalian cells for structural studies. Nat. Protoc. 9, 2574–2585. https://doi.org/10.1038/nprot.2014.173

Goldschen-Ohm, M.P., Haroldson, A., Jones, M.V., Pearce, R.A., 2014. A nonequilibrium binary elements-based kinetic model for benzodiazepine regulation of GABAA receptors. J. Gen. Physiol. 144, 27–39. https://doi.org/10.1085/jgp.201411183

Grant, T., Rohou, A., Grigorieff, N., 2018. cisTEM, user-friendly software for single-particle image processing. eLife 7. https://doi.org/10.7554/eLife.35383

Harrison, N.J., Lummis, S.C.R., 2006. Locating the Carboxylate Group of GABA in the Homomeric rho GABAA Receptor Ligand-binding Pocket*. J. Biol. Chem. 281, 24455–24461. https://doi.org/10.1074/jbc.M601775200

Hill, D.R., Bowery, N.G., 1981. 3H-baclofen and 3H-GABA bind to bicuculline-insensitive GABAB sites in rat brain. Nature 290, 149–152. https://doi.org/10.1038/290149a0

Huang, J., Rauscher, S., Nawrocki, G., Ran, T., Feig, M., de Groot, B.L., Grubmüller, H., MacKerell, A.D.J., 2017. CHARMM36m: an improved force field for folded and intrinsically disordered proteins. Nat. Methods 14, 71–73. https://doi.org/10.1038/nmeth.4067

Ingólfsson, H.I., Carpenter, T.S., Bhatia, H., Bremer, P.-T., Marrink, S.J., Lightstone, F.C., 2017. Computational Lipidomics of the Neuronal Plasma Membrane. Biophys. J. 113, 2271–2280. https://doi.org/10.1016/j.bpj.2017.10.017

Jansen, M., Bali, M., Akabas, M.H., 2008. Modular Design of Cys-loop Ligand-gated Ion Channels: Functional 5-HT3 and GABA ρ1 Receptors Lacking the Large Cytoplasmic M3M4 Loop. J. Gen. Physiol. 131, 137–146. https://doi.org/10.1085/jgp.200709896

Jianliang Zhang, Fenqin Xue, Yongchang Chang, 2008. Structural Determinants for Antagonist Pharmacology That Distinguish the ρ_1_ GABA_C_ Receptor from GABA_A_ Receptors. Mol. Pharmacol. 74, 941. https://doi.org/10.1124/mol.108.048710

Jo, S., Kim, T., Iyer, V.G., Im, W., 2008. CHARMM-GUI: a web-based graphical user interface for CHARMM. J. Comput. Chem. 29, 1859–1865. https://doi.org/10.1002/jcc.20945

Jones, S.M., Palmer, M.J., 2009. Activation of the Tonic GABAC Receptor Current in Retinal Bipolar Cell Terminals by Nonvesicular GABA Release. J. Neurophysiol. 102, 691–699. https://doi.org/10.1152/jn.00285.2009

Jorgensen, W.L., Chandrasekhar, J., Madura, J.D., Impey, R.W., Klein, M.L., 1983. Comparison of simple potential functions for simulating liquid water. J. Chem. Phys. 79, 926–935. https://doi.org/10.1063/1.445869

Jumper, J., Evans, R., Pritzel, A., Green, T., Figurnov, M., Ronneberger, O., Tunyasuvunakool, K., Bates, R., Žídek, A., Potapenko, A., Bridgland, A., Meyer, C., Kohl, S.A.A., Ballard, A.J., Cowie, A., Romera-Paredes, B., Nikolov, S., Jain, R., Adler, J., Back, T., Petersen, S., Reiman, D., Clancy, E., Zielinski, M., Steinegger, M., Pacholska, M., Berghammer, T., Bodenstein, S., Silver, D., Vinyals, O., Senior, A.W., Kavukcuoglu, K., Kohli, P., Hassabis, D., 2021. Highly accurate protein structure prediction with AlphaFold. Nature 596, 583–589. https://doi.org/10.1038/s41586-021-03819-2

Kash, T.L., Jenkins, A., Kelley, J.C., Trudell, J.R., Harrison, N.L., 2003. Coupling of agonist binding to channel gating in the GABA(A) receptor. Nature 421, 272–275. https://doi.org/10.1038/nature01280

Khatib, F., Cooper, S., Tyka, M.D., Xu, K., Makedon, I., Popović, Z., Baker, D., Players, F., 2011. Algorithm discovery by protein folding game players. Proc. Natl. Acad. Sci. 108, 18949–18953. https://doi.org/10.1073/pnas.1115898108

Kim, J.J., Gharpure, A., Teng, J., Zhuang, Y., Howard, R.J., Zhu, S., Noviello, C.M., Walsh, R.M., Lindahl, E., Hibbs, R.E., 2020. Shared structural mechanisms of general anaesthetics and benzodiazepines. Nature 585, 303–308. https://doi.org/10.1038/s41586-020-2654-5

Kim, J.J., Hibbs, R.E., 2021. Direct Structural Insights into GABA(A) Receptor Pharmacology. Trends Biochem. Sci. 46, 502–517. https://doi.org/10.1016/j.tibs.2021.01.011

Kimanius, D., Dong, L., Sharov, G., Nakane, T., Scheres, S.H.W., 2021. New tools for automated cryo-EM single-particle analysis in RELION-4.0. Biochem. J. 478, 4169–4185. https://doi.org/10.1042/BCJ20210708

Krissinel, E., Henrick, K., 2007. Inference of macromolecular assemblies from crystalline state. J. Mol. Biol. 372, 774–797. https://doi.org/10.1016/j.jmb.2007.05.022

Kumar, A., Basak, S., Rao, S., Gicheru, Y., Mayer, M.L., Sansom, M.S.P., Chakrapani, S., 2020. Mechanisms of activation and desensitization of full-length glycine receptor in lipid nanodiscs. Nat. Commun. 11, 3752. https://doi.org/10.1038/s41467-020-17364-5

Laverty, D., Desai, R., Uchański, T., Masiulis, S., Stec, W.J., Malinauskas, T., Zivanov, J., Pardon, E., Steyaert, J., Miller, K.W., Aricescu, A.R., 2019. Cryo-EM structure of the human α1β3γ2 GABAA receptor in a lipid bilayer. Nature 565, 516–520. https://doi.org/10.1038/s41586-018-0833-4

Licari, G., Dehghani-Ghahnaviyeh, S., Tajkhorshid, E., 2022. Membrane Mixer: A Toolkit for Efficient Shuffling of Lipids in Heterogeneous Biological Membranes. J. Chem. Inf. Model. 62, 986–996. https://doi.org/10.1021/acs.jcim.1c01388

Liu, H., Fu, H., Chipot, C., Shao, X., Cai, W., 2021. Accuracy of Alternate Nonpolarizable Force Fields for the Determination of Protein-Ligand Binding Affinities Dominated by Cation-π Interactions. J. Chem. Theory Comput. 17, 3908–3915. https://doi.org/10.1021/acs.jctc.1c00219

Lyons, J.A., Bøggild, A., Nissen, P., Frauenfeld, J., 2017. Saposin-Lipoprotein Scaffolds for Structure Determination of Membrane Transporters. Methods Enzymol. 594, 85–99. https://doi.org/10.1016/bs.mie.2017.06.035

Masiulis, S., Desai, R., Uchański, T., Serna Martin, I., Laverty, D., Karia, D., Malinauskas, T., Zivanov, J., Pardon, E., Kotecha, A., Steyaert, J., Miller, K.W., Aricescu, A.R., 2019. GABAA receptor signalling mechanisms revealed by structural pharmacology. Nature 565, 454–459. https://doi.org/10.1038/s41586-018-0832-5

Matthews, G., Ayoub, G., Heidelberger, R., 1994. Presynaptic inhibition by GABA is mediated via two distinct GABA receptors with novel pharmacology. J. Neurosci. 14, 1079. https://doi.org/10.1523/JNEUROSCI.14-03-01079.1994

Miller, P.S., Aricescu, A.R., 2014. Crystal structure of a human GABAA receptor. Nature 512, 270–275. https://doi.org/10.1038/nature13293

Mirdita, M., Schütze, K., Moriwaki, Y., Heo, L., Ovchinnikov, S., Steinegger, M., 2022. ColabFold: making protein folding accessible to all. Nat. Methods 19, 679–682. https://doi.org/10.1038/s41592-022-01488-1

Mirdita, M., Steinegger, M., Söding, J., 2019. MMseqs2 desktop and local web server app for fast, interactive sequence searches. Bioinforma. Oxf. Engl. 35, 2856–2858. https://doi.org/10.1093/bioinformatics/bty1057

Morales-Perez, C.L., Noviello, C.M., Hibbs, R.E., 2016. Manipulation of Subunit Stoichiometry in Heteromeric Membrane Proteins. Struct. Lond. Engl. 1993 24, 797–805. https://doi.org/10.1016/j.str.2016.03.004

Murata, Y., Woodward, R.M., Miledi, R., Overman, L.E., 1996. The first selective antagonist for a GABAC receptor. Bioorg. Med. Chem. Lett. 6, 2073–2076. https://doi.org/10.1016/0960-894X(96)00364-2

Naffaa, M.M., Absalom, N., Solomon, V.R., Chebib, M., Hibbs, D.E., Hanrahan, J.R., 2016. Investigating the Role of Loop C Hydrophilic Residue ‘T244’ in the Binding Site of ρ1 GABAC Receptors via Site Mutation and Partial Agonism. PLOS ONE 11, e0156618. https://doi.org/10.1371/journal.pone.0156618

Naffaa, M.M., Hung, S., Chebib, M., Johnston, G.A.R., Hanrahan, J.R., 2017. GABA-ρ receptors: distinctive functions and molecular pharmacology. Br. J. Pharmacol. 174, 1881–1894. https://doi.org/10.1111/bph.13768

Noviello, C.M., Gharpure, A., Mukhtasimova, N., Cabuco, R., Baxter, L., Borek, D., Sine, S.M., Hibbs, R.E., 2021. Structure and gating mechanism of the α7 nicotinic acetylcholine receptor. Cell 184, 2121–2134.e13. https://doi.org/10.1016/j.cell.2021.02.049

Páll, S., Zhmurov, A., Bauer, P., Abraham, M., Lundborg, M., Gray, A., Hess, B., Lindahl, E., 2020. Heterogeneous parallelization and acceleration of molecular dynamics simulations in GROMACS. J. Chem. Phys. 153, 134110. https://doi.org/10.1063/5.0018516

Parrinello, M., Rahman, A., 1980. Crystal Structure and Pair Potentials: A Molecular-Dynamics Study. Phys. Rev. Lett. 45, 1196–1199. https://doi.org/10.1103/PhysRevLett.45.1196

Pédelacq, J.-D., Cabantous, S., Tran, T., Terwilliger, T.C., Waldo, G.S., 2006. Engineering and characterization of a superfolder green fluorescent protein. Nat. Biotechnol. 24, 79–88. https://doi.org/10.1038/nbt1172

Petroff, J.T., Dietzen, N.M., Santiago-McRae, E., Deng, B., Washington, M.S., Chen, L.J., Trent Moreland, K., Deng, Z., Rau, M., Fitzpatrick, J.A.J., Yuan, P., Joseph, T.T., Hénin, J., Brannigan, G., Cheng, W.W.L., 2022. Open-channel structure of a pentameric ligand-gated ion channel reveals a mechanism of leaflet-specific phospholipid modulation. Nat. Commun. 13, 7017. https://doi.org/10.1038/s41467-022-34813-5

Pettersen, E.F., Goddard, T.D., Huang, C.C., Meng, E.C., Couch, G.S., Croll, T.I., Morris, J.H., Ferrin, T.E., 2021. UCSF ChimeraX: Structure visualization for researchers, educators, and developers. Protein Sci. Publ. Protein Soc. 30, 70–82. https://doi.org/10.1002/pro.3943

Phulera, S., Zhu, H., Yu, J., Claxton, D.P., Yoder, N., Yoshioka, C., Gouaux, E., 2018. Cryo-EM structure of the benzodiazepine-sensitive α1β1γ2S tri-heteromeric GABAA receptor in complex with GABA. eLife 7, e39383. https://doi.org/10.7554/eLife.39383

Polenzani, L., Woodward, R.M., Miledi, R., 1991. Expression of mammalian gamma-aminobutyric acid receptors with distinct pharmacology in Xenopus oocytes. Proc. Natl. Acad. Sci. 88, 4318–4322. https://doi.org/10.1073/pnas.88.10.4318

Price, K.L., Millen, K.S., Lummis, S.C.R., 2007. Transducing Agonist Binding to Channel Gating Involves Different Interactions in 5-HT3 and GABAC Receptors *. J. Biol. Chem. 282, 25623–25630. https://doi.org/10.1074/jbc.M702524200

Punjani, A., Zhang, H., Fleet, D.J., 2020. Non-uniform refinement: adaptive regularization improves single-particle cryo-EM reconstruction. Nat. Methods 17, 1214–1221. https://doi.org/10.1038/s41592-020-00990-8

Qian, H., Dowling, J.E., Ripps, H., 1998. Molecular and pharmacological properties of GABA-ρ subunits from white perch retina. J. Neurobiol. 37, 305–320. https://doi.org/10.1002/(SICI)1097-4695(19981105)37:2<305::AID-NEU9>3.0.CO;2-6

Qian, H., Pan, Y., Zhu, Y., Khalili, P., 2005. Picrotoxin Accelerates Relaxation of GABA_C_ Receptors. Mol. Pharmacol. 67, 470. https://doi.org/10.1124/mol.104.003996

Schmidt, T.G.M., Batz, L., Bonet, L., Carl, U., Holzapfel, G., Kiem, K., Matulewicz, K., Niermeier, D., Schuchardt, I., Stanar, K., 2013. Development of the Twin-Strep-tag® and its application for purification of recombinant proteins from cell culture supernatants. Protein Expr. Purif. 92, 54–61. https://doi.org/10.1016/j.pep.2013.08.021

Smart, O.S., Neduvelil, J.G., Wang, X., Wallace, B.A., Sansom, M.S.P., 1996. HOLE: A program for the analysis of the pore dimensions of ion channel structural models. J. Mol. Graph. 14, 354–360. https://doi.org/10.1016/S0263-7855(97)00009-X

Smart O.S., Sharff, A., Holstein, J., Womack, T.O., Flensburg, C., Keller, P., Paciorek, W., Vonrhein, C., Bricogne, G., 2021. Grade, version 1.2.20.

String method solution of the gating pathways for a pentameric ligand-gated ion channel., 2017. United States. https://doi.org/10.1073/pnas.1617567114

Vanommeslaeghe, K., Hatcher, E., Acharya, C., Kundu, S., Zhong, S., Shim, J., Darian, E., Guvench, O., Lopes, P., Vorobyov, I., Mackerell, A.D.J., 2010. CHARMM general force field: A force field for drug-like molecules compatible with the CHARMM all-atom additive biological force fields. J. Comput. Chem. 31, 671–690. https://doi.org/10.1002/jcc.21367

Vanommeslaeghe, K., MacKerell, A.D.J., 2012. Automation of the CHARMM General Force Field (CGenFF) I: bond perception and atom typing. J. Chem. Inf. Model. 52, 3144–3154. https://doi.org/10.1021/ci300363c

Vanommeslaeghe, K., Raman, E.P., MacKerell, A.D.J., 2012. Automation of the CHARMM General Force Field (CGenFF) II: assignment of bonded parameters and partial atomic charges. J. Chem. Inf. Model. 52, 3155–3168. https://doi.org/10.1021/ci3003649

Vikram Dalal, Mark J. Arcario, John T. Petroff II, Noah M. Dietzen, Michael J. Rau, James A. J. Fitzpatrick, Grace Brannigan, Wayland W. L. Cheng, 2022. Lipid nanodisc scaffold and size alters the structure of a pentameric ligand-gated ion channel. bioRxiv 2022.11.20.517256. https://doi.org/10.1101/2022.11.20.517256

Wallace, A.C., Laskowski, R.A., Thornton, J.M., 1995. LIGPLOT: a program to generate schematic diagrams of protein-ligand interactions. Protein Eng. Des. Sel. 8, 127–134. https://doi.org/10.1093/protein/8.2.127

Walters, R.J., Hadley, S.H., Morris, K.D.W., Amin, J., 2000. Benzodiazepines act on GABAA receptors via two distinct and separable mechanisms. Nat. Neurosci. 3, 1274–1281. https://doi.org/10.1038/81800

Wang, T.L., Hackam, A.S., Guggino, W.B., Cutting, G.R., 1995. A single amino acid in gamma-aminobutyric acid rho 1 receptors affects competitive and noncompetitive components of picrotoxin inhibition. Proc. Natl. Acad. Sci. 92, 11751–11755. https://doi.org/10.1073/pnas.92.25.11751

Williams, C.J., Headd, J.J., Moriarty, N.W., Prisant, M.G., Videau, L.L., Deis, L.N., Verma, V., Keedy, D.A., Hintze, B.J., Chen, V.B., Jain, S., Lewis, S.M., Arendall, W.B. 3rd, Snoeyink, J., Adams, P.D., Lovell, S.C., Richardson, J.S., Richardson, D.C., 2018. MolProbity: More and better reference data for improved all-atom structure validation. Protein Sci. Publ. Protein Soc. 27, 293–315. https://doi.org/10.1002/pro.3330

Wotring, V.E., Chang, Y., Weiss, D.S., 1999. Permeability and single channel conductance of human homomeric ρ1 GABAC receptors. J. Physiol. 521, 327–336. https://doi.org/10.1111/j.1469-7793.1999.00327.x

Xiu, X., Hanek, A.P., Wang, J., Lester, H.A., Dougherty, D.A., 2005. A unified view of the role of electrostatic interactions in modulating the gating of Cys loop receptors. J. Biol. Chem. 280, 41655–41666. https://doi.org/10.1074/jbc.M508635200

Yang, L., Nakayama, Y., Hattori, N., Liu, B., Inagaki, C., 2008. GABAC-Receptor Stimulation Activates cAMP-Dependent Protein Kinase via A-Kinase Anchoring Protein 220. J. Pharmacol. Sci. 106, 578–584. https://doi.org/10.1254/jphs.FP0071362

Yang, L., Omori, Kyoko, Omori, Koichiro, Otani, H., Suzukawa, J., Inagaki, C., 2003. GABAC receptor agonist suppressed ammonia-induced apoptosis in cultured rat hippocampal neurons by restoring phosphorylated BAD level. J. Neurochem. 87, 791–800. https://doi.org/10.1046/j.1471-4159.2003.02069.x

Yoshikawa, N., Hutchison, G.R., 2019. Fast, efficient fragment-based coordinate generation for Open Babel. J. Cheminformatics 11, 49. https://doi.org/10.1186/s13321-019-0372-5

Yu, J., Zhu, H., Lape, R., Greiner, T., Du, J., Lü, W., Sivilotti, L., Gouaux, E., 2021. Mechanism of gating and partial agonist action in the glycine receptor. Cell 184, 957–968.e21. https://doi.org/10.1016/j.cell.2021.01.026

Zhu, F., Feng, M., Sinha, R., Murphy, M.P., Luo, F., Kao, K.S., Szade, K., Seita, J., Weissman, I.L., 2019. The GABA receptor GABRR1 is expressed on and functional in hematopoietic stem cells and megakaryocyte progenitors. Proc. Natl. Acad. Sci. 116, 18416–18422. https://doi.org/10.1073/pnas.1906251116

Zhu, S., Noviello, C.M., Teng, J., Walsh, R.M., Kim, J.J., Hibbs, R.E., 2018. Structure of a human synaptic GABAA receptor. Nature 559, 67–72. https://doi.org/10.1038/s41586-018-0255-3

